# A genomic snapshot of demographic and cultural dynamism in Upper Mesopotamia during the Neolithic Transition

**DOI:** 10.1101/2022.01.31.478487

**Authors:** N. Ezgi Altınışık, Duygu Deniz Kazancı, Ayça Aydoğan, Hasan Can Gemici, Ömür Dilek Erdal, Savaş Sarıaltun, Kıvılcım Başak Vural, Dilek Koptekin, Kanat Gürün, Ekin Sağlıcan, Gökhan Çakan, Meliha Melis Koruyucu, Vendela Kempe Lagerholm, Cansu Karamurat, Mustafa Özkan, Gülşah Merve Kılınç, Arda Sevkar, Elif Sürer, Anders Götherström, Çiğdem Atakuman, Yılmaz Selim Erdal, Füsun Özer, Aslı Erim Özdoğan, Mehmet Somel

## Abstract

Upper Mesopotamia played a key role in the Neolithic Transition in Southwest Asia through marked innovations in symbolism, technology, and foodways. We present thirteen ancient genomes (c.8500-7500 calBCE) from Pre-Pottery Neolithic Çayönü in the Tigris basin together with bioarchaeological and material culture data. Our findings reveal that Çayönü was a genetically diverse population, carrying a mixed ancestry from western and eastern Fertile Crescent, and that the community received immigrants. Our results further suggest that the community was organised along biological family lines. We document bodily interventions such as head-shaping and cauterization among the individuals examined, reflecting Çayönü’s cultural ingenuity. Finally, we identify Upper Mesopotamia as the likely source of eastern gene flow into Neolithic Anatolia, in line with material culture evidence. We hypothesise that Upper Mesopotamia’s cultural dynamism during the Neolithic Transition was the product not only of its fertile lands but also of its interregional demographic connections.

## Introduction

Located between the Euphrates and Tigris rivers, the hilly flanks of Upper Mesopotamia were home to the earliest sedentary hunter-gatherers who built the first monumental structures at Göbekli Tepe (*1*) and domesticated numerous local plant and animal species, including einkorn, emmer, sheep, goat, pig, and cattle (*2–5*). The innovative spirit and cultural dynamism of these societies during the Neolithic Transition in Southwest Asia (c. 9800-6500 BC) is well documented in the archaeological record, but their demographic history and social structures has remained unknown owing to the lack of genomes from North Mesopotamia. This stands in contrast with a significant number of recent archaeogenomic studies that focused on the three most distant corners of Neolithic Southwest Asia, namely South Levant, Central Zagros, and Central Anatolia (Fig. 1A, B) (*6–12*). This body of work has together revealed (a) genetically distinct populations in all three regions, (b) a dominant trend of population continuity between pre-Neolithic, Pre-Pottery Neolithic (PPN) and Pottery Neolithic (PN) communities, (c) an overlay of interregional gene flow through time, such as inferred “southern” and “eastern” gene flow events into Central Anatolia between the Early and Late Neolithic. Meanwhile, key questions about the possible roles of Upper Mesopotamia in interregional demographic and cultural change, e.g., whether Upper Mesopotamia influenced Late Neolithic Central Anatolia and whether it was the source of the post-Neolithic gene flow into Anatolia (*6, 13*), have remained open. With the exception of a single ancient DNA study reporting 15 mitochondrial DNA sequences from the Upper Euphrates (*14*), Upper Mesopotamia has remained genomically unexplored, mostly owing to low DNA preservation in the region.

**Fig. 1.**
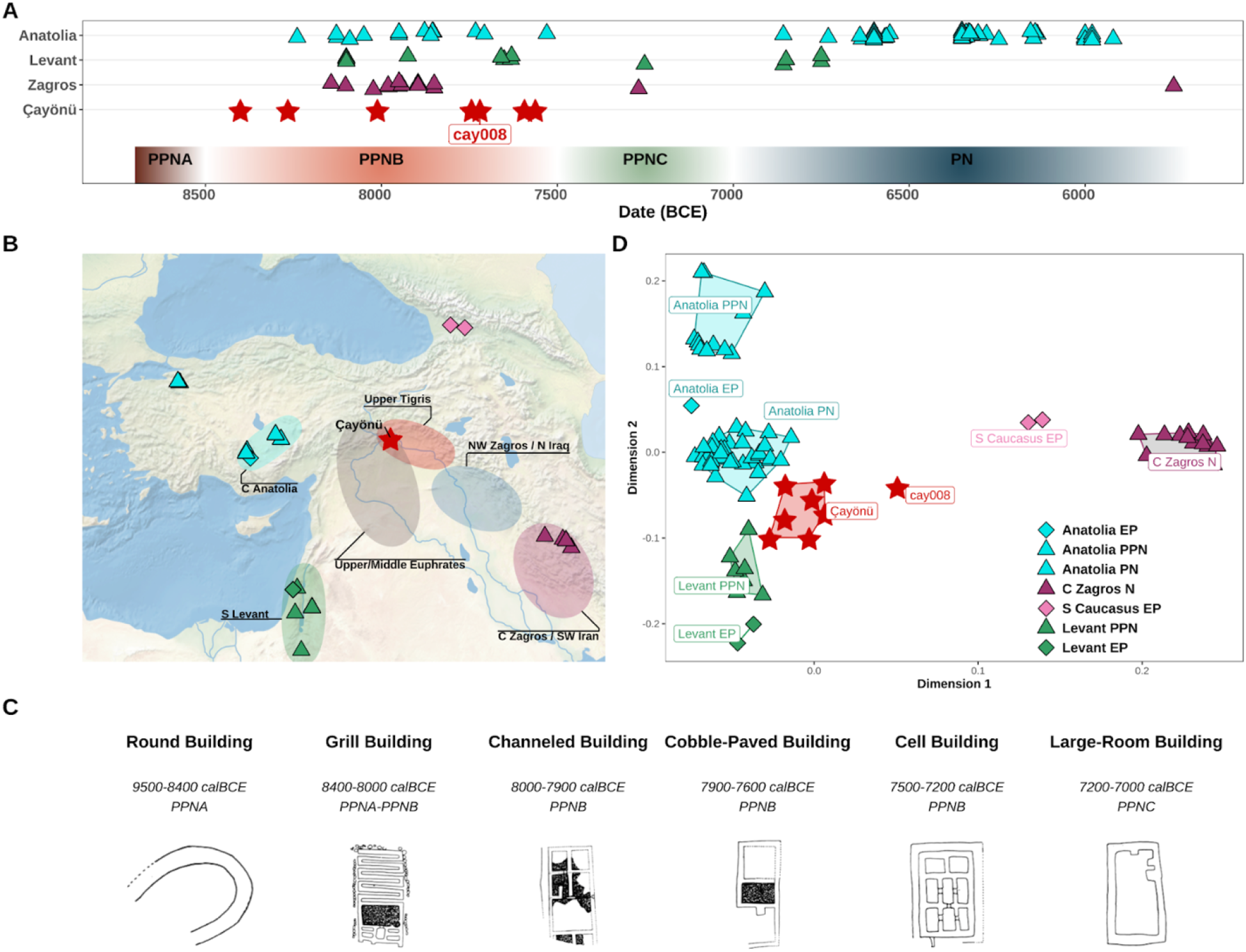
Spatio-temporal distribution of the samples and the population structure of Neolithic Southwest Asia. **(A)** Timeline of ancient Southwest Asian individuals used in the analyses. Colored rectangles at the bottom represent the sub-periods of the Neolithic Era in Southwest Asia. **(B)** The map shows Epi-Paleolithic (EP) and Neolithic populations from Southwest Asia. Shaded areas mark Pre-Pottery Neolithic (PPN) cultural zones. **(C)** Çayönü building types and their approximate dates of use, considered as evidence for Çayönü’s cultural openness and ingenuity. Adapted from (*103*). **(D)** First two dimensions of multidimensional scaling (MDS) plot of genetic distances. The MDS summarises the genetic distance matrix among ancient genomes calculated as (1 - outgroup *f*_*3*_) values. Outgroup *f*_3_-statistics were calculated as *f*_3_*(Yoruba; individual_1_, individual_2_)*. The labels represent the following sites: Anatolia EP: Pınarbaşı; Anatolia PPN: Boncuklu and Aşıklı Höyük; Anatolia PN: Çatalhöyük and Barcın Höyük; Levant EP: Natufian; Levant PPN: Ain’ Ghazal, Kfar HaHoresh, Motza and Ba’ja; C Zagros N (Central Zagros Neolithic): Ganj Dareh, Tepe Abdul and Wezmeh Cave; S Caucasus EP (South Caucasus EP): Kotias and Satsurblia. We use “Anatolia” here following the traditional geographic definition, referring to the west of the Anatolian Diagonal.

Here we address this gap by studying genomic data from Çayönü Tepesi (hereon Çayönü) of the Upper Tigris area (Fig. 1A), a settlement that presents one of the best examples of the transition from foraging to food production in Southwest Asia (*15*). Firstly, Çayönü’s uninterrupted stratigraphy extending from the Pre-Pottery Neolithic A (PPNA) (c. 9500 cal BCE) to the final PPN (c. 7000 cal BCE) is unparalleled in the region. Secondly, the Çayönü Neolithic community is recognized for its marked cultural dynamism, which is reflected (a) in evidence for intense plant cultivation (*16*) and animal management (pig, cattle, sheep and goat) (*17*), (b) in continuous innovation in architectural styles (Fig. 1C), (c) in technological experimentation, from pioneering lime burning techniques (*15, 18*) to the production of copper beads and reamer-like objects (*19*). Finally, both western (Levant-Euphrates) and eastern (Tigris-Zagros) influences and parallel developments are traceable in Çayönü’s material culture (*20*) (Supplementary Table 1). These observations suggest that Çayönü and contemporaneous Upper Mesopotamian communities could have acted as hubs of cultural interaction and innovation in Neolithic Southwest Asia.

Our study presents genomic data from Çayönü, which we then use to describe (i) the structure of Fertile Crescent populations in comparison with interregional material culture affinities, (ii) the Neolithic demographic transition reflected in intraregional genomic diversity, (iii) genetic kinship among co-burials in domestic structures at Çayönü, and (iv) the potential role of Upper Mesopotamia in Neolithic and post-Neolithic human movements influencing Anatolia. We also detail the curious case of a Çayönü infant, whom we infer to be a migrant offspring, and whose skeletal material presents the earliest known examples of cauterization and head-shaping in the region.

## Results and Discussion

We studied a total of 33 human remains from Çayönü (Fig. 1A, B, Supplementary Table 2). These were mainly found as subfloor burials located inside or within the proximity of six Pre-Pottery Neolithic B (PPNB) buildings (Table S1). We screened 33 aDNA libraries by shotgun sequencing, which revealed endogenous DNA proportions varying between 0.04% and 5% (median = 0.2%, Fig. S1). This was lower than aDNA preservation in a contemporaneous Central Anatolian settlement, Aşıklı (median = 1.4%, Wilcoxon rank sum test p < 0.05), but comparable to another Central Anatolian site, Boncuklu (median = 0.1%, Wilcoxon rank sum test p > 0.05, Fig. S2).

Libraries from 14 individuals were chosen for deeper sequencing (Methods), from which we generated shotgun genomes with depths ranging from 0.016x to 0.49x. (Fig. S1, Supplementary Table 2). High rates of post-mortem damage (PMD) accumulation at read ends, short average fragment sizes (49-60 bps, median = 51.4 bps), and mitochondrial haplotype-based estimates suggested authenticity of all 14 libraries (Methods) (Supplementary Table 2). With this data we first estimated genetic kinship among all individual pairs (Methods). Two samples, both identified as female infants (cay018 and cay020), were genetically inferred to either belong to the same individual or to be identical twins. Anthropological evaluation also showed that both petrouses could belong to the same individual. We therefore merged their genomic data and treated this merged data as representing a single individual, reducing our sample size to 13 individuals (6 adult females, 2 adult males, 3 sub-adult females, 2 sub-adult males). We further identified four related pairs of 1^st^ to 3^rd^ degree (see below) and removed all but one individual among sets of closely related individuals in population genetic analyses (Methods).

### The east-west genetic structure of Neolithic Southwest Asia

To obtain an overview of genetic affinities among human populations in Neolithic Southwest Asia we compared the 13 Çayönü genomes with published ancient genomes dating to c.15,000-5,500 BCE from the Fertile Crescent and neighbouring regions (Supplementary Table 3) (*6–8, 11, 12, 21–24*) using multidimensional scaling (MDS) of pairwise *f_3_* results, *D*-statistics, and *qpAdm* analyses (*25*). These revealed a number of observations. In the MDS analysis, the Çayönü group occupied a distinct and intermediate position within the space of Southwest Asian genetic diversity bordered by early Holocene South Levant, Central Zagros and South Caucasus, and Central Anatolia (Fig. 1D, Fig. S3). Our sample of Çayönü genomes was internally homogeneous within this space, with the exception of an “outlier” individual, cay008, who appeared relatively closer to Zagros/Caucasus individuals. D-statistics likewise showed that the Çayönü group was genetically closer to western Southwest Asia (early Holocene Central Anatolia and South Levant) than to eastern Southwest Asia (Central Zagros) (Fig. 2A, Supplementary Table 4). At the same time, Central Zagros genomes showed higher genetic affinity to our Çayönü sample than to Central Anatolia or South Levant (Fig. 2B). Finally, we found that cay008 harbours higher Zagros contribution than other Çayönü individuals (Fig. 1D, Fig. 2C, Supplementary Table 5).

**Fig. 2.**
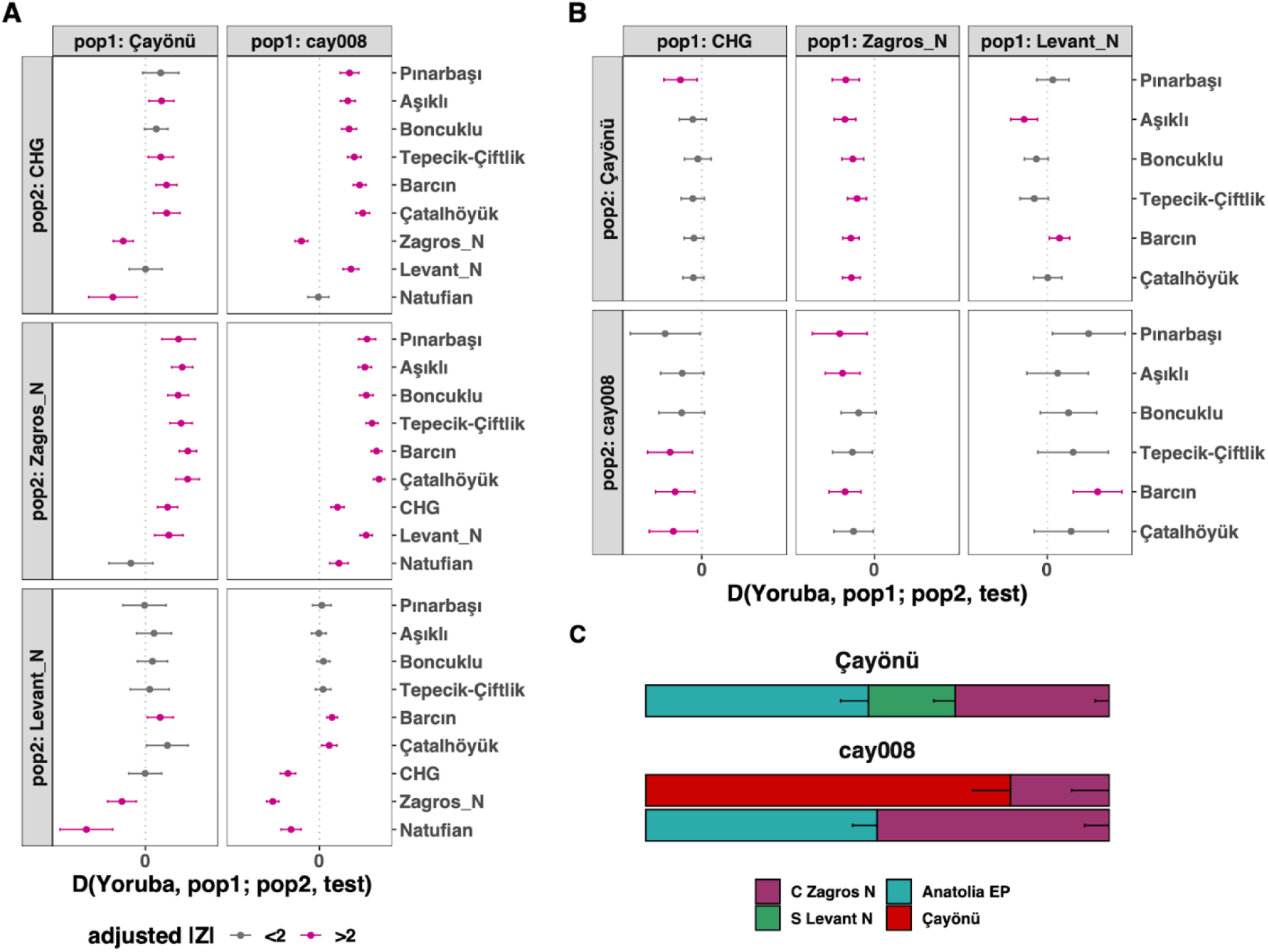
Genetic affinities of Çayönü population with the neighbouring populations. Formal tests computed in the form of **(A)** *D(Yoruba, Çayönü/cay008; pop2, test) D(Yoruba, pop1; Çayönü/cay008, Anatolia EP/PPN/PN)*. Z-scores were corrected with the Benjamini-Hochberg multiple testing correction (*82*). Horizontal bars represent ±2 standard errors. **(C)** *qpAdm* modelling of the Çayönü group and cay008. The “local” Çayönü group or outlier cay008 individual was the “target”; Central Anatolia Epi-Paleolithic, Central Zagros Neolithic and South Levant Neolithic samples were sources for both targets. The “local” Çayönü group was also used as “source” for modelling of cay008. Horizontal bars represent standard errors of the coefficients. All three models yielded *p*-values > 0.05. We also cannot reject a three-way model of Central Anatolia PPN, Central Zagros and South Levant at >0.01 *p*-value threshold (Supplemental Table 5). In all analyses shown in the figure, “Çayönü” represents the 9 genomes listed in Table 1, excluding relatives and cay008.

Given these observations, we first investigated the origins of the genetic structure in Neolithic Southwest Asia. The higher genetic affinity among Upper Mesopotamia (represented by Çayönü), Central Anatolia, and South Levant populations relative to Central Zagros (Fig. 1D, Fig. S3) was intriguing, which led us to ask whether this affinity could be explained by an isolation-by-distance process (*26, 27*). We computed shared genetic drift between each pair of individuals in our Southwest Asia sample and compared these to geodesic geographic distance among settlements (Methods). To eliminate the effect of temporal genetic changes, we only included individual pairs separated by <1,000 years. We found a general correlation between spatial and genetic distances, as expected (Fig. 3A). However, we also found that Central Zagros genomes were significantly more differentiated compared to that expected from a linear isolation-by-distance model (Fig. 3B). We therefore infer a stark west-east genetic structure in the Fertile Crescent, where the lowest effective migration (*28*) appears to lie between Upper Mesopotamia and Central Zagros.

**Fig. 3.**
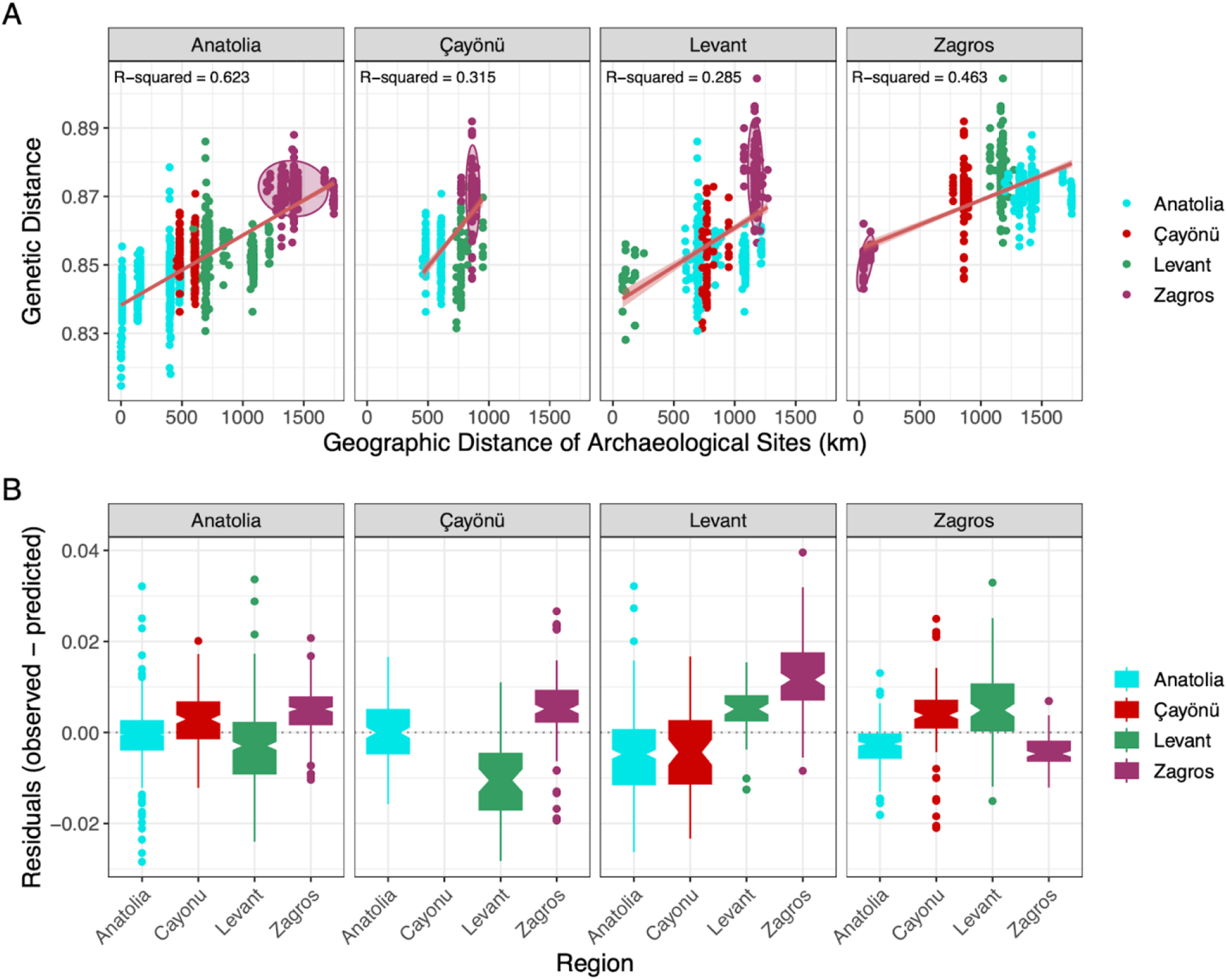
Genetic isolation-by-distance in Southwest Asia. Panel **(A)** shows correlation between geographic (x-axis) and genetic (y-axis) distance for Southwest Asia Neolithic populations. The red regression line shows the linear fit with 95% confidence interval. Each point represents pairs of individuals from Southwest Asia Neolithic. Pairs from the same site, and pairs separated by >1,000 years time difference were not included. All regression lines were highly significant (*p*<0.001). Panel **(B)** shows the distribution of residuals which we calculated by subtracting observed values from predicted values obtained from the linear regression models in Panel A. In all analyses shown in the figure, “Çayönü” represents the 9 genomes listed in Table 1, excluding relatives and cay008.

At face value, this result may seem to imply resistance to gene flow between Upper Mesopotamia and Central Zagros during the Neolithic. However, such resistance does not align with observed material culture affinities between the two regions (e.g., (*29*) Supplementary Table 1). We therefore suggest an alternative scenario to explain the observed genetic structure. During Last Glacial Maximum (LGM), the eastern regions of Southwest Asia (the ancestors of Central Zagros / South Caucasus) could have been isolated from the western regions (the ancestors of Central Anatolians/Levantines), with the east and west populations differentiating through drift or by admixture with third populations. Sometime after the LGM, these eastern and western regional populations could have re-expanded and partly admixed within Southwest Asia. It is plausible that east-west admixture occurred in Upper Mesopotamia, giving rise to Çayönü’s gene pool, and may have also influenced Central Anatolia by the PN (*6*). We note that the duration and timing of this putative admixture process remain unclear, and that alternative scenarios are also conceivable (*30*). Irrespective of the demographic mechanisms, though, Central Zagros appears to have been genetically the most distinct group in early Holocene Southwest Asia.

### Admixed ancestry and diverse material culture affinities in Çayönü

We next investigated the demographic origins of Çayönü inhabitants. The *D*-statistics results mentioned above had suggested that the Çayönü sample carried mixed eastern and western ancestry (Fig. 2A, B), which is consistent with the site’s intermediate geographic position. Using *qpAdm* we could further model ancestry proportions in the Çayönü genome sample (excluding the cay008 individual) as three-way admixtures of Central Anatolia-(represented by Epi-Paleolithic (EP) Pınarbaşı; Anatolia_EP), South Levant- and Central Zagros-related ancestries (Fig. 2C; Supplemental Table 5) (Methods). Çayönü bears mainly Anatolian ancestry, complemented by 33% (SE ± 3%) of Zagros and 19% (SE ± 5%) of Southern Levant ancestry (p-value > 0.05).

We then asked whether the genetic affinity of Çayönü individuals to regional populations could have changed over the 1,000 years covered by our sample (Fig. 4). We found no significant temporal effect (multiple testing corrected p>0.05). Still, this does not rule out immigration into Çayönü, as the cay008 “outlier” individual shows. With *qpAdm* we estimated that the cay008 genome carried 50% Anatolia_EP and 50% Zagros_N ancestry (SE ± 5%, p-value > 0.05) and lacked a significant South Levant component found in the rest of Çayönü genomes (Fig. 2C). We were also able to model cay008 as a mixture of the “local” Çayönü sample (79%, SE ± 8%) and Zagros-like (21%, SE ± 8%) ancestries (p-value > 0.05). We caution that “Zagros ancestry” here might actually represent Northwest Zagros ancestry (i.e., modern-day North Iraq, from where archaeogenomic data is not yet available) rather than Central Zagros ancestry, which would also be compatible with Çayönü’s material cultural affinities with Northwest Zagros (Fig. 1B; Supplementary Table 1).

**Fig. 4.**
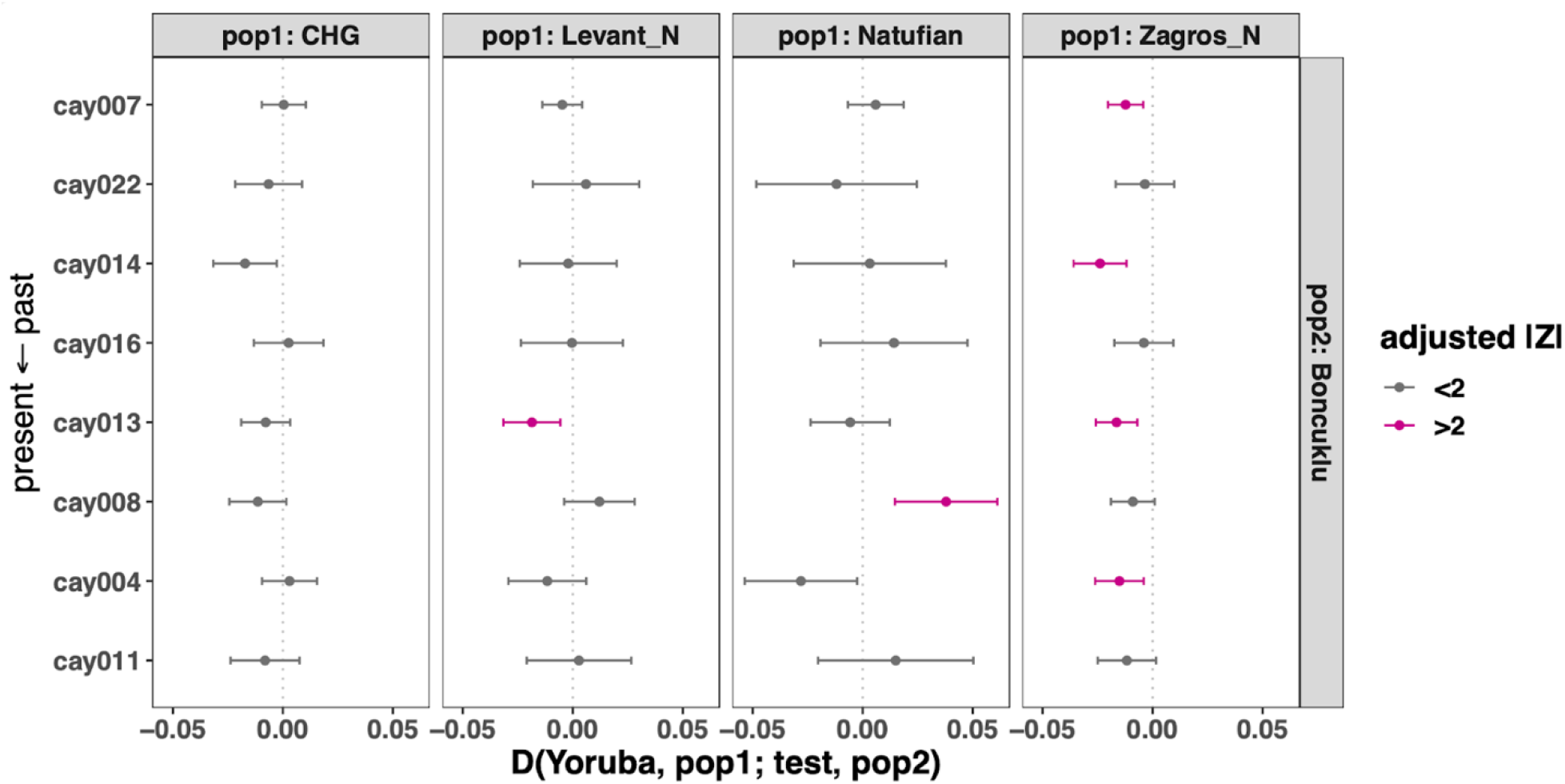
Testing temporal genetic change in Çayönü. Formal tests computed in the form of *D*(*Yoruba, pop1; test, Boncuklu*) where *pop1* denotes CHG/Levant_N/Natufian/Zagros_N and *test* denotes radiocarbon-dated Çayönü individuals (Table 1), ordered from past to present in top to bottom direction, respectively. Horizontal bars represent ±2 standard errors.

These observations overall support the notion that the Çayönü population had both historical and ongoing demographic connections with the neighbouring regions. Archaeologically, Çayönü shares a number of distinctive features with PPNA/PPNB settlements in the eastern wing of Neolithic Southwest Asia, particularly those in the Tigris and Euphrates basins and Northwest Zagros (Fig. 1B; Supplementary Table 1). These features include monumental architecture and/or special buildings, lithic types such as the “Çayönü tool”, and plain or winged marble bracelets (*15, 20, 31, 32*) (Supplementary Table 1). Another observation worth mentioning is the joint presence of both the pressure technique and bidirectional blade technologies at Çayönü, which were predominant in the eastern and western regions of Neolithic Southwest Asia, respectively (Supplementary Table 1). Obsidian network analyses further suggest close interactions between the Tigris and Zagros areas (*29*). We speculate that Çayönü’s east-west mixed ancestry and its possible openness to interregional human movement may have facilitated its observed wide-ranging material culture affinities and cultural dynamism.

### A pre-agriculture demographic shift in the “Fertile Crescent”

Our dataset further allowed us to revisit a previous observation on the demographic impact of the Neolithic transition. It had been earlier reported that the Central Anatolian PPN (pre-agriculturalist) populations Aşıklı and Boncuklu had low levels of genetic diversity, similar to Upper Paleolithic and Mesolithic Europeans and Caucasians (*8, 12*). In comparison, Central Anatolian PN populations Tepecik-Çiftlik and Çatalhöyük, as well as West Anatolian and European Neolithic populations carried higher genetic diversity levels. This temporal increase in genetic diversity was attributed to the transition to farming and the consequent intensification of population movements and admixture ((*8*); also see (*11*)).

Here we asked whether PPN populations of the “Fertile Crescent” *sensu stricto*, which comprises the main domestication centres of animals and plants in Southwest Asia, also had low genetic diversity levels similar to those of Central Anatolian PPN groups. Measuring genetic diversity using outgroup *f_3_* values in genetic samples from Upper Mesopotamia (Çayönü), South Levant (Ain Ghazal) and Central Zagros (Ganj Dareh), we found that genetic samples from all three “Fertile Crescent” PPN settlements had high diversity levels, on a par with later-coming agriculturalists of Central and Western Anatolia PN, and significantly higher than those of Central Anatolia PPN (Fig. 5A, Supplementary Table 6).

**Fig. 5.**
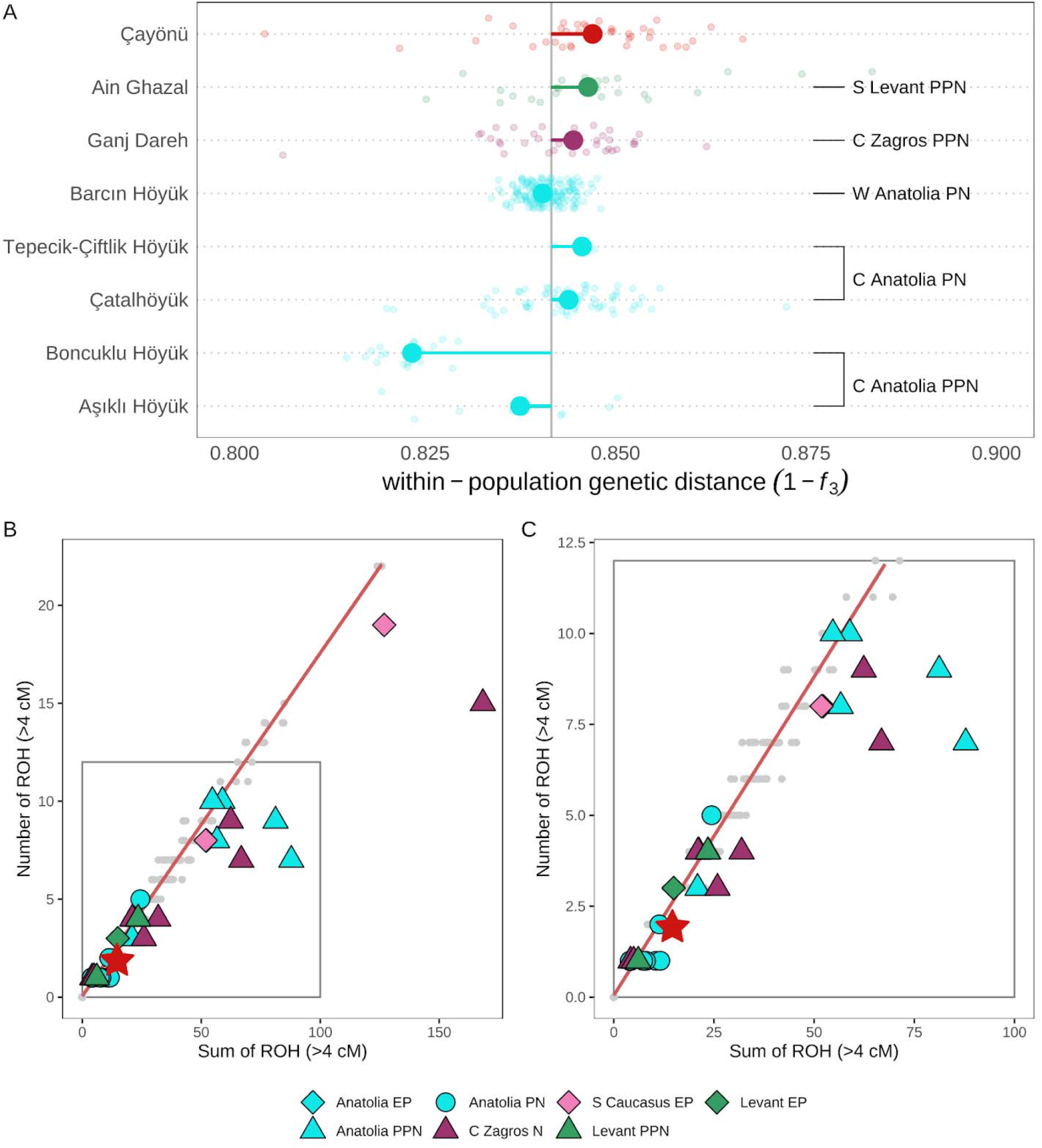
Genetic diversity in Neolithic Southwest Asia. **(A)** Small dots show pairwise genetic distance calculated as (1 - outgroup *f*_*3*_) values for all pairs of individuals, whereas large dots show the median values of each population. The vertical grey line represents the total median across the 8 populations. Deviation from the total median is shown with colored horizontal lines. Outgroup *f*_*3*_-statistics were computed as *f_3_(Yoruba; ind_1_, ind_2_)* where *ind_1_* and *ind_2_* represent individuals from the same archaeological site. **(B)** and **(C)** presents runs of homozygosity (ROH) in Southwest Asia. Sum of total ROH >4 cM and Number of total ROH >4 cM was shown on the x- and y-axes, respectively. The baseline (the red diagonal line) was computed using short ROH values (4-8 cM) in present-day West and Central Eurasian individuals to represent outbreed samples to determine the baseline. Panel is the zoomed version of Panel **(B)** in which we draw the zoomed area with grey rectangular. Red star point is the only Çayönü individual, namely cay007, that have more than 300,000 SNPs in the 1240K SNP Panel. The grey dots designate the ROH values for modern genomes.

We further studied background population diversity through runs of homozygosity (ROH) analyses of one Çayönü genome (cay007) with sufficient coverage using the *hapROH* algorithm (*33*) and compared these with ROH distributions estimated in other early Holocene Southwest Asian genomes (Methods). As reported earlier (*8, 12, 21, 23*), Aşıklı, Boncuklu, and Caucasus pre-Neolithic genomes carried large numbers of ROH, indicative of small population size. Certain Neolithic genomes (e.g., WC1, Ash128) also showed a “right-shift” when plotting the number versus sum of ROHs, indicative of recent inbreeding (*34*) (Fig. 5B-C). In contrast, the cay007 genome had small and few ROHs, suggesting lack of recent inbreeding and a relatively large population size, respectively.

Overall, these results suggest that the demographic transition observed in Central Anatolia between the PPN and the PN did not take place in the “Fertile Crescent”, at least at the same magnitude. This observation is in line with radiocarbon-based estimates of low population density on the Central Anatolian plateau relative to other Fertile Crescent regions in the early Holocene (*35*). This result would also be consistent with the “Hilly Flanks hypothesis”, according to which the fertile flanks of the Taurus and Zagros Mountains that supported both the large hunter-gatherer populations and the progenitors of plant and animal domesticates eventually became cradles of early domestication events (*36–38*).

### Çayönü co-burials reflect nuclear and extended family structures

Recent work suggested that in the Central Anatolian PPN communities, Aşıklı Höyük and Boncuklu Höyük, co-burials frequently included close genetic kin, suggesting that these earliest sedentary communities may have been organised around biological families (*12*), as hypothesised earlier (*39, 40*). In contrast, co-burials were rarely close kin in PN communities, Çatalhöyük and Barcın (*12*), implying distinct social organisation patterns in the latter. But the generality of these observations has remained uncertain as this single study included limited sample sizes. Here we investigated genetic kinship among Çayönü co-burials using two approaches. First, using methods that have sensitivity up to second- or third-degree relatedness (*41, 42*), we estimated genetic kinship levels among a total of 78 pairs, with 10 pairs representing co-burials in the same building. We identified 4 closely related pairs, including first-, second-, and third-degree relationships (Fig. 6A-B, Supplementary Table 7-8). These 4 pairs were interred in three buildings, and each pair shared the same building (Fig. 7).

**Fig. 6.**
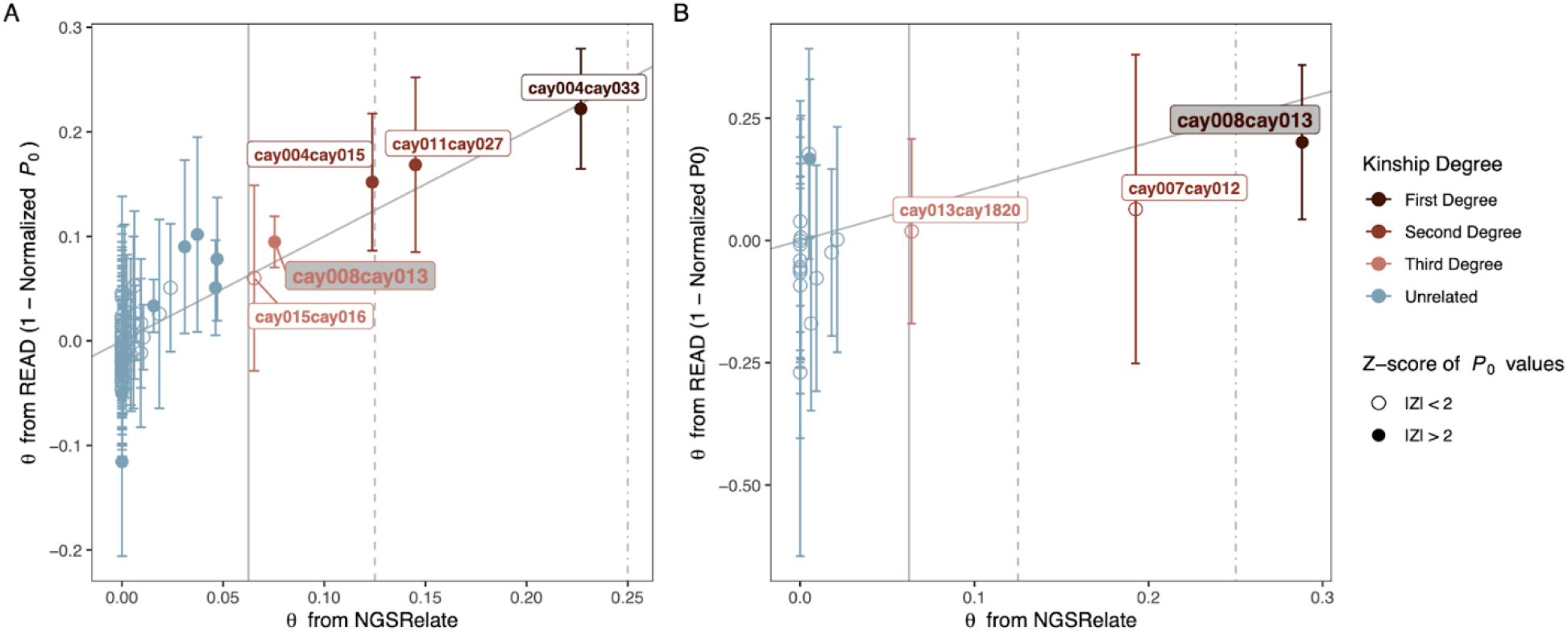
Kinship coefficient (θ) estimates among Çayönü individuals. Comparison of **(A)** autosomal and **(B)** X-chromosomal estimates of θ are shown. In both panels, *NGSRelate* θ estimates are shown on the x-axes and *READ* θ estimates, calculated as (1 - normalised *P_0_*), on the y-axes. Vertical bars represent ±2 standard errors of *P_0_* values. Vertical dotted, dashed and straight grey lines intersect with expected θ values for first-, second- and third-degree relatives, respectively. Annotation with the grey label shows the pair cay008 and cay013.

**Fig. 7.**
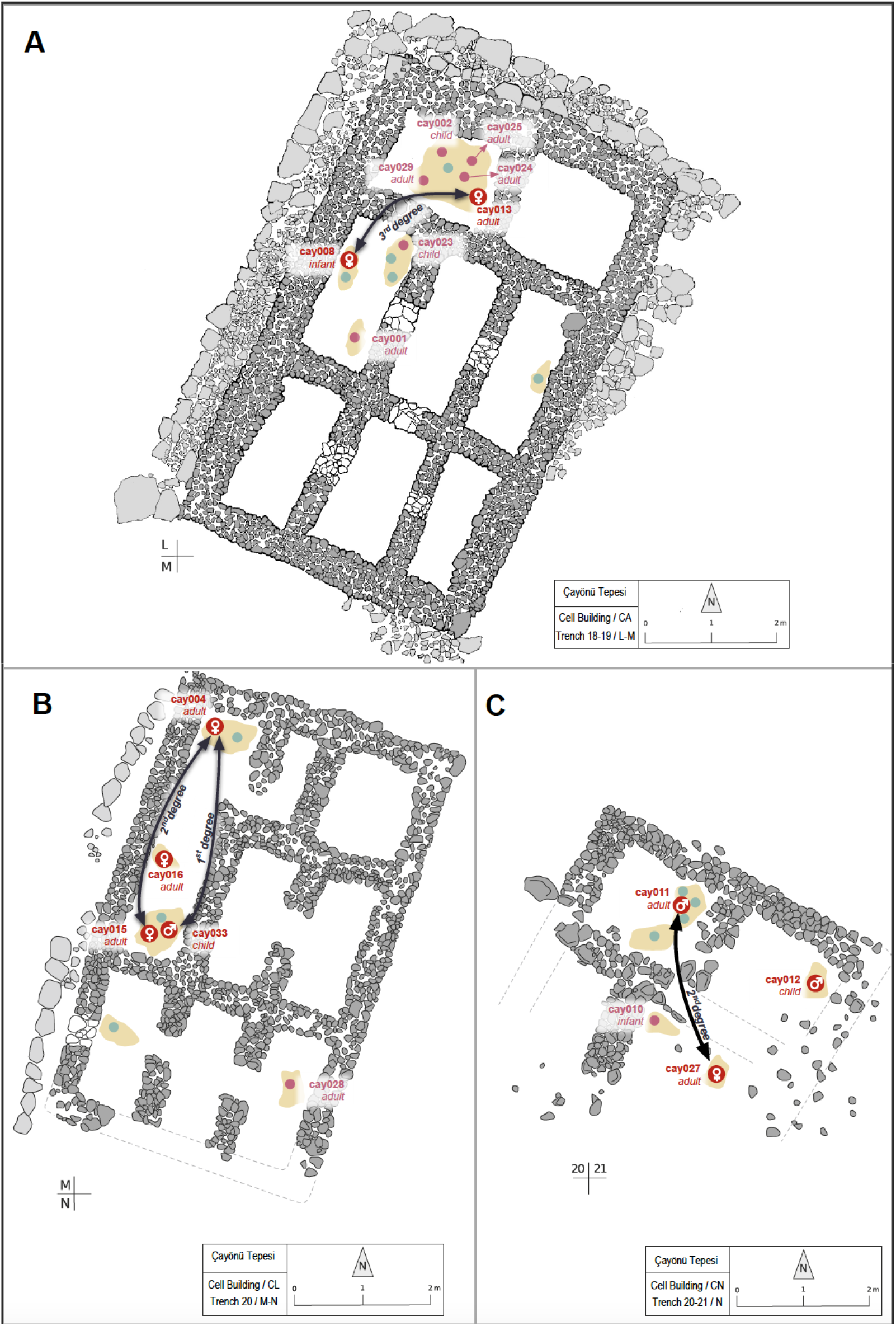
Locations of Çayönü co-burials interred in domestic buildings. All three buildings belong to the Cell Building sub-phase. The figure shows plans of buildings coded **(A)** CA, **(B)** CL, and **(C)** CN. Red dots represent individuals analysed in this study; pink dots represent individuals screened for aDNA but with insufficient preservation, and blue dots represent burials of other individuals within the same buildings. Black curved lines show the closely related pairs in each building.

We hypothesised that the 9 individuals who were co-buried with others but were not closely related, could still belong to the same extended biological families. We investigated this by testing whether each of these co-buried pairs were genetically closer to each other than to other Çayönü individuals, using outgroup *f_3_*-statistics. We indeed found that co-buried pairs who were not identified as close genetic kin were still slightly genetically closer to each other than pairs from distinct buildings (effect size = 0.03, permutation test *p-*value < 0.001) (Fig. S4).

Our results are similar to observations from the contemporaneous Aşıklı and Boncuklu of Central Anatolian PPN (*12*), but not genetic kinship analyses of Upper Palaeolithic co-burials from Sunghir, Siberia (*43*). Hence, biological family-based co-burial cultures may belong to a diverse set of derived cultural traits shared among communities across Southwest Asia in this period. The social significance of Neolithic co-burials and whether they represented household members is yet ambiguous, though the observed patterns are consistent with the notion that biological family structures played a role in social organisation in Southwest Asia during the transition to sedentism (*39, 40*). Our results also render the reported deficiency of a genetic kinship signal in the later-coming Barcın and Çatalhöyük PN communities (*44, 45*) even more intriguing.

### A toddler of migrant descent, with intentional head-shaping and cauterization

Our genetic comparisons highlighted a 1.5-2-year-old female toddler, cay008, as an outlier, with higher genetic affinity to Zagros populations (Fig. 1D, Fig. 2, Fig. S3). Genetic kinship analysis using autosomal loci suggested a third-degree relationship between this individual and an adult female, cay013, interred in the same building (Fig. 6–7, Supplementary Table 7). In contrast, analysis of their X-chromosomal loci indicated a genetic relationship closer than third-degree (Fig. 6B, Supplementary Table 8). Such discrepancy would be expected if cay013 was the paternal kin of cay008. Pedigree analysis suggested cay013 being the paternal great-aunt of cay008 as a likely scenario (Methods; Fig. S5 and 6). In addition, the mtDNA haplogroup of cay008 (haplogroup T2g) was a clear outlier within the Çayönü sample, which consisted mostly of haplogroup K1 (Supplementary Table 2). These lines of evidence suggest that the Zagros-like ancestry of cay008 was inherited from her maternal side and her migrant ancestors bred with local individuals.

This toddler further displayed two intriguing features in her cranium (Fig. 8A-F). First, cay008’s skull appears to be subject to intentional head-shaping, manifested as frontal flattening with a fronto-occipital groove and post-coronal depression (Fig. 8C). This could be produced by a double-bandaged circular head-shaping procedure. Three additional individuals in our sample also showed similar evidence, including cay013, the adult female relative of cay008 (Table S1). Although circular head-shaping with two bandages was previously documented in Neolithic Southwest Asia (*46, 47*), Çayönü presents one of the earliest known examples of this tradition.

**Fig. 8.**
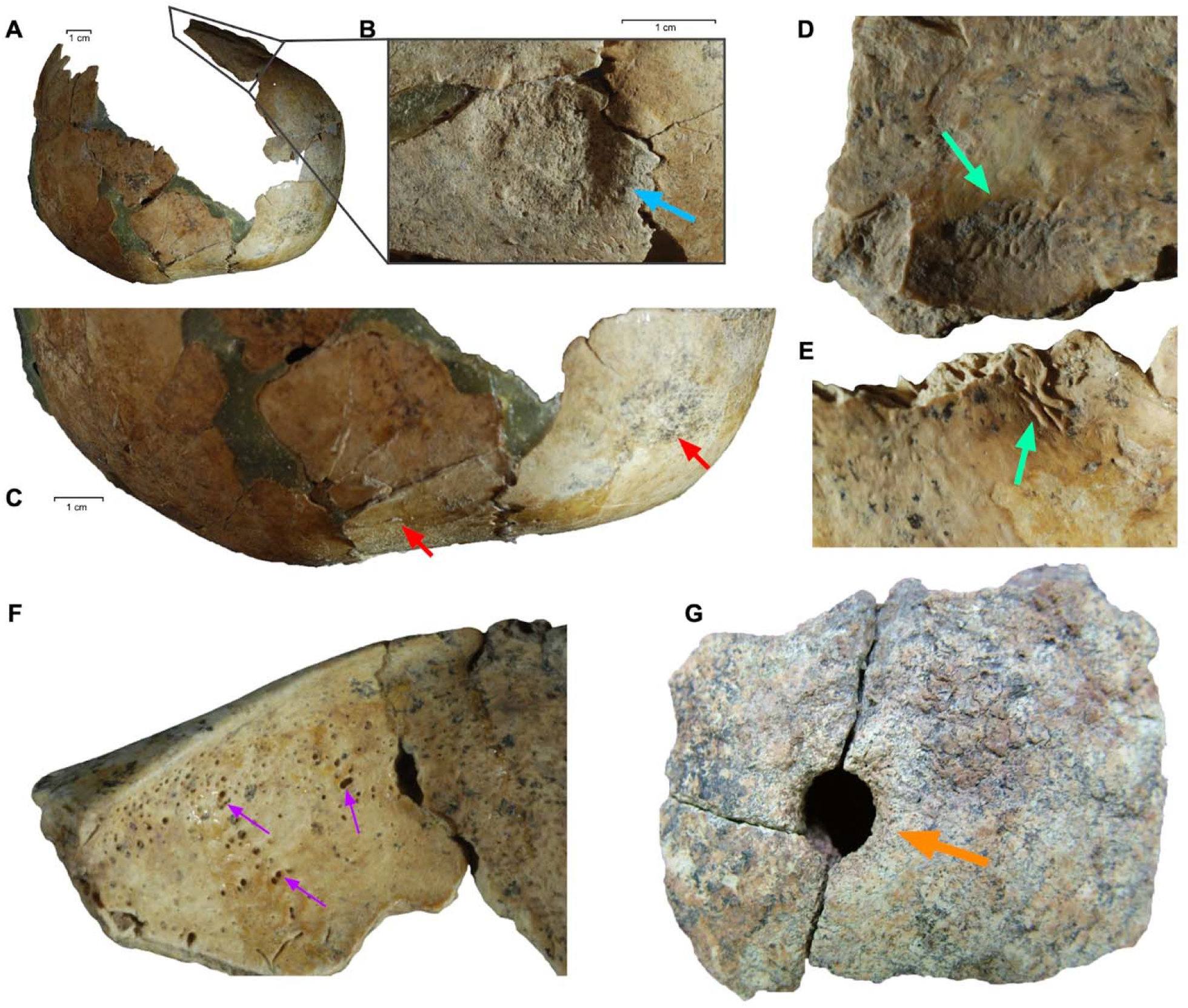
Cranial features of the cay008 toddler. **(A)** Frontal flattening, post-coronal depression, bulging on the parietal tuber and fronto-occipital grooving suggest a double bandaged circular type cranial deformation. **(B)** Cauterisation with a circular depression found on the post-coronal area on the left parietal bone. The bone is very thin in the centre and the edge of the lesion is elevated. **(C)** An enlarged picture of a post-coronal depression and frontal flattening. **(D-E)** Endocranial lithic lesions similar to *serpens endocrania symmetrica* on occipital bone. **(F)** Slightly developed cribra orbitalia on the right orbital roof. This lesion together with porotic hyperostosis is mainly related to anaemia. **(G)** Cranial trepanation performed by drilling on the skull of Çayönü individual ÇT’78 KE 6-2/3a SK5 (not represented in our genetic sample).

The cay008 skull also presents evidence of cauterization, i.e., the intentional burning of the cranium by an instrument (Fig. 8B). Cauterization marks were prevalent in Neolithic populations in Anatolia and Europe (*47, 48*), but to our knowledge, cay008 shows the earliest documented case of this treatment. Cauterization marks from Europe are usually associated with trepanation, performed to thin the cranial bone (*49*), but the cranium of cay008 lacks a trepanation signal. Instead, we observed endocranial lesions reminiscent of *serpens endocrania symmetrica* on the inner surface of the fragmented occipital of cay008, suggesting that the toddler suffered from an infection (Fig. 8D, E). The cranium also showed *cribra orbitalia* which can signal anaemia (Fig. 8F). We hypothesise that cauterization on the parietal bone might have been applied to treat the adverse effect of these diseases. The bone formation suggests the toddler lived for a period of time after cauterization.

The evidence for cauterization, widespread head shaping, and additional reports of trepanation in Çayönü (Fig. 8G) altogether suggest a prominent culture of intentional body modification in this community (*50*). Body modifications may have developed in parallel with other aspects of cultural innovation in Çayönü, and could also be shared interregionally; indeed, cases of head-shaping and trepanation are also known from Neolithic sites in the Fertile Crescent (*46, 51–53*).

### The demographic impact of Upper Mesopotamia on Neolithic and post-Neolithic Anatolia

Finally, we investigated the possible role of Upper Mesopotamia as a source of post-7,000 BCE eastern gene flow into Anatolia. Eastern gene flow events have been inferred from increasing levels of early Holocene South Caucasus and/or Zagros ancestry in Anatolian populations, starting by the PN and continuing into the Bronze Age (*8, 13, 54, 55*). It was speculated that the original source of Caucasus/Zagros-related ancestry might be Upper Mesopotamia (*13, 55*). This could be plausible given our results above, specifically that our Çayönü sample included >25% Zagros ancestry relative to Central Anatolian PPN populations (Fig. 2C). We thus asked whether the post-7,000 BCE eastern admixture in Anatolian populations is better explained by gene flow from an early Holocene Caucasus-related group, or from Upper Mesopotamia, represented by Çayönü. We computed *D*-statistics in the form of *D(Yoruba, CHG/Çayönü; X, Anatolia_EP/Anatolia_PPN)*, where *X* was a Neolithic to Bronze Age population from Anatolia/Aegean, *CHG* represents early Holocene Caucasus (the so-called “Caucasus hunter-gatherers”), and *Anatolia_EP/Anatolia_PPN* represents Epi-Palaeolithic Pınarbaşı and Central Anatolia PPN Boncuklu, respectively.

This revealed two interesting results. First, we found that Çayönü genomes show higher genetic affinity to PN Anatolian populations Çatalhöyük, Tepecik-Çiftlik, and Barcın, than to pre-7000 BCE Anatolian genomes (Fig. 9A, B; Supplementary Table 4). Moreover, this affinity was weak or absent when using CHG instead of Çayönü (*D*-statistics ≈ 0). This result is consistent with Upper Mesopotamia, but not Caucasus, being the source of eastern gene flow into Central Anatolia and possibly Western Anatolia around 7,000 BCE. The finding also resonates with archaeological evidence from Çatalhöyük, where the mid-7th millennium BCE witnesses the first introduction of obsidian from the Bingöl area of Eastern Turkey, the appearance of lithic types akin to “Çayönü tools”, and an increasing use of the pressure technique in lithic industries (*56–58*).

**Fig. 9.**
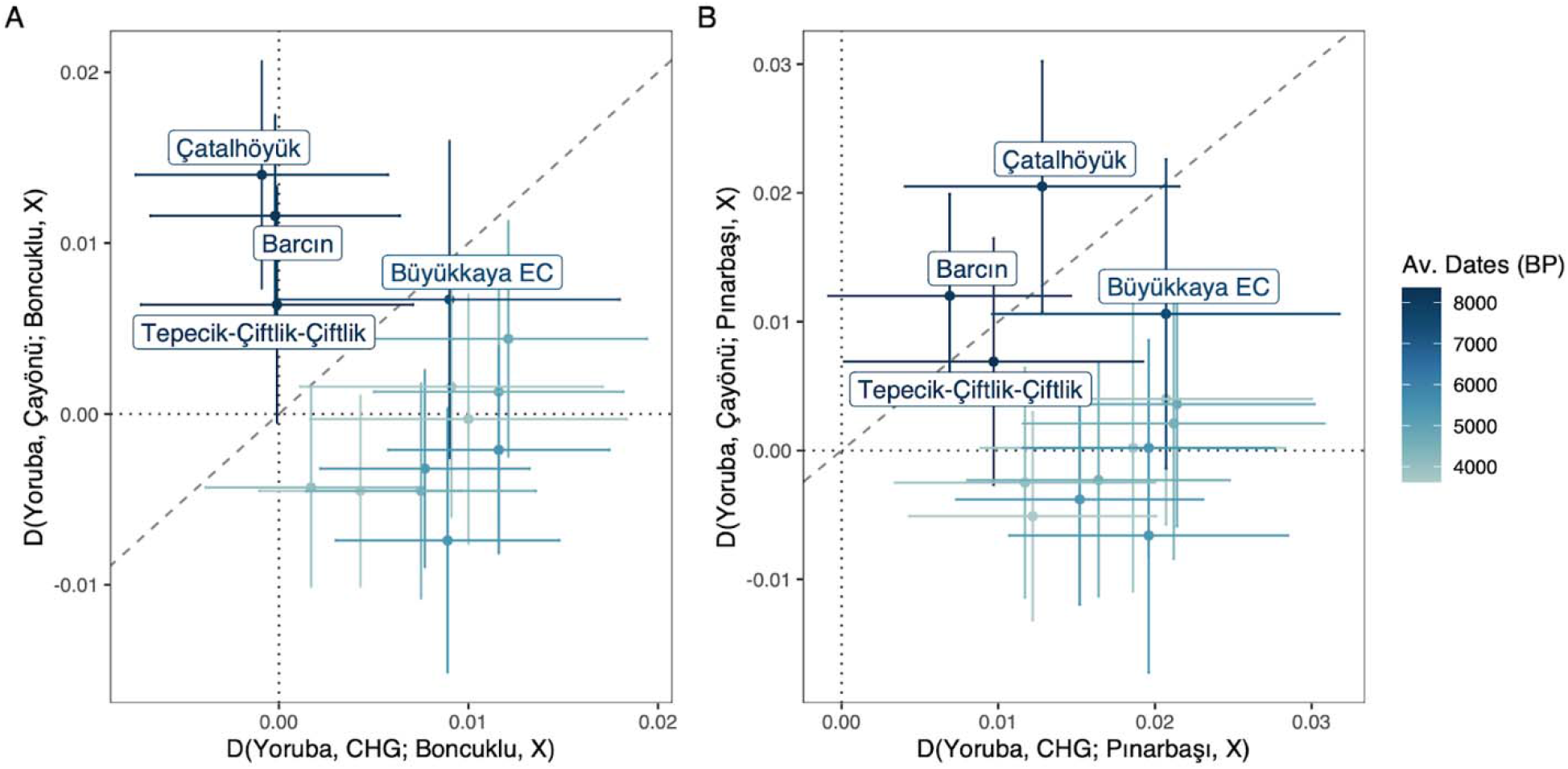
Biplots of *D*-statistics illustrating excess allele sharing between Çayönü and post-7000 BCE populations from Central/Western Anatolia. *D*-statistics were computed in the form of *D(Yoruba, pop1; pop2, X)* where *X* represents Pottery Neolithic, Chalcolithic and Bronze Age populations from the Anatolian Plateau. Each population is represented by a dot and error bars representing ±2 standard errors. The list of populations and *D*-statistics can be found in Supplementary Table 4. In both panels, *pop1* corresponds to CHG on the x-axes whereas on the y-axes *pop1* corresponds to the Çayönü population. *pop2* is represented by Boncuklu (Central Anatolian PPN) in panel **(A)** and Pınarbaşı (Central Anatolian EP) in panel **(B)** in both axes. The slope of the diagonal dashed line is 1 showing x = y and the intercept of both vertical and horizontal dotted lines are 0.

Second, starting with early Chalcolithic in Anatolia, Çayönü genomes lose their affinity to post-Neolithic Anatolians while the CHG sample gains affinity to post-Neolithic Anatolians over pre-7000 BCE Anatolians (Fig. 9A-B). Hence, PPN Çayönü-related groups do not appear as the direct source of Caucasus-related ancestry in post-Neolithic Anatolia. This can be explained in two ways. One is that Caucasus-related influence in post-Neolithic populations emerged from another region than Upper Mesopotamia, such as North or East Anatolia. An alternative scenario is that the Upper Mesopotamian gene pool itself changed post-7000 BCE by Zagros/Caucasus-related gene flow. In this case, Upper Mesopotamia could have remained the source of eastern gene flow into Anatolia with its new profile.

## Conclusion

Whereas the main driver behind European Neolithization has been recognized as mass population movements from Anatolia and/or Southeast Europe (*9, 22, 24*), the role of human movement in the multi-millennia process of Neolithization in Southwest Asia is less understood. Here, we described the formation of Upper Mesopotamian PPN populations, represented by PPNB Çayönü, as an admixture event between western and eastern populations of early Holocene Southwest Asia. The PPNB Çayönü community appears to have carried relatively high genetic diversity levels relative to PPN Central Anatolia and pre-Neolithic Europe which indicates that the site was open to interaction.

Nearly half a century ago archaeologists Robert Braidwood and Halet Çambel described Çayönü as a perfect spot for the emergence of sedentism and agriculture, due its location along the Hilly Flanks of the Taurus and Zagros Mountains where progenitors of plant and domesticates naturally co-existed (*59*). We hypothesise that Çayönü was also a lively hub of interregional networks, potentially due to its location between the sources of the Tigris and the Euphrates rivers in Upper Mesopotamia. Recent discoveries and ongoing research at sites such as Göbekli Tepe and Karahan Tepe (*60, 61*) continue to demonstrate the significance of this region as a central node of cultural dynamism and social networks.

## Materials and Methods

### Laboratory procedures

#### Sample collection and direct radiocarbon dating

Petrous bones from 33 human individuals from Çayönü, housed at the Hacettepe University in Ankara, Turkey, were used in aDNA experiments (Fig. 1A, Table 1). Table S1 provides archaeological and anthropological background information of all individuals. Eight of the deep sequenced samples were dated with the AMS C14 method at TUBITAK-MAM (Gebze, Turkey). Radiocarbon ages were calibrated using the *IntCal13* calibration curve (*62*) with *OxCal* v4.2 (*63*).

**Table 1.** Description of sequenced Çayönü individuals. (*) denotes individuals used in population genetic analyses to represent the “local” Çayönü population.

| Excavation ID | C14 dates (cal BC) | Sample ID | Average read length | Genome coverage (X) | Genetic sex | Note |
| --- | --- | --- | --- | --- | --- | --- |
| ÇT’78 S.16 | 7649-7538 | cay004 | 51.44 | 0.08 | XX | * |
| ÇT’81 S.2a | 8496-8306 | cay007 | 54.40 | 0.49 | XY | * |
| ÇT’86 S.2 | 7842-7598 | cay008 | 51.51 | 0.08 | XX | Identified as genetic “outlier” |
| ÇT’78 S.7 | 7601-7524 | cay011 | 52.56 | 0.04 | Consistent with XY but not XX | * |
| ÇT’78 S.6 | - | cay012 | 52.03 | 0.04 | XY | * |
| ÇT’78 S.21 | 7882-7606 | cay013 | 49.92 | 0.14 | XX | * |
| ÇT’81 HB isolated | 8211-7812 | cay014 | 50.74 | 0.04 | XX | * |
| ÇT’81 S.15 | - | cay015 | 51.40 | 0.02 | XX | Relative of cay004 |
| ÇT’81 S.8 | 7882-7606 | cay016 | 52.31 | 0.04 | XX | * |
| ÇT’78 S.25 | 8300-8232 | cay022 | 60.96 | 0.04 | XX | * |
| ÇT’78 S.1 | - | cay027 | 49.63 | 0.02 | XX | Relative of cay011 |
| ÇT’84 S.60 | - | cay033 | 51.89 | 0.02 | Consistent with XY but not XX | Relative of cay004 |
| ÇT’70 S.13/<br>ÇT’70 S.11b | - | cay1820 | 50.89 | 0.07 | XX | *<br>Identical individuals/ Merged libraries |

#### Ancient DNA extraction, whole genome library preparation and sequencing

The experiments were carried out in dedicated ancient DNA facilities at the Middle East Technical University and Hacettepe University in Ankara, Turkey. In order to prevent contamination, all equipment and utensils were decontaminated with DNA AWAY or a bleach solution at each use and also in between handling the samples. UV-insensitive solutions were UV irradiated for 10 minutes at a distance of 5 cm prior to use. Negative controls were included at each step of the experiments to be able to track potential contamination originating from the reagents or handling the samples.

Prior to DNA extraction, outer surfaces of the petrous bones were scraped to a depth of c.1 millimetre with single-use scalpels. The cochlea and the surrounding compact bone were cut out using the Dremel tool (*64*) and a piece from this region was ground into a fine powder using a SPEX 6770 freezer mill. About 120 mg of bone powder was transferred to a 2 ml screwtop tube. Ancient DNA extraction was performed following the Dabney et al. (*65*) protocol. Two tubes of 80 mg hydroxyapatite, one at the beginning, the other at the end of each set of extractions, were used as negative controls. Blunt-end, double-stranded, Illumina compatible sequencing libraries with double-indexes were prepared using the Kircher et al. (*66*) protocol and sequenced on the Illumina Novaseq 6000 platform using Novaseq S1 flowcells at low coverage (median c. 26 million reads per sample) (Supplementary Table 2). After alignment (see the “Sequencing Data Preprocessing” Section below), 14 individuals’ libraries were found to contain >0.3% endogenous DNA; these were further sequenced on the Illumina Novaseq 6000 platform using S1 flowcells.

### Quantification and statistical analysis

#### Sequencing data preprocessing

We removed Illumina adaptor sequences in *fastq* files and merged the paired-end sequencing reads using *AdapterRemoval* v2.3.1 (*67*), requiring an overlap of at least 11 bps between pairs. The merged reads were mapped against the Human Reference Genome (hs37d5), using the *“samse”* command of *BWA aln* v0.7.15 (*68*) with the parameters “*-n 0.01 -o 2*” and with seeding disabled using the “*−l 16500*” option. PCR duplicates were removed using the *FilterUniqueSAMCons.py* script by collapsing the reads with identical start and end positions (*69*). Finally, we filtered reads shorter than 35 bps length and with more than 10% mismatches to the reference genome. Multiple libraries belonging to the same individual were merged using *SAMtools merge* v1.9 (*68*), and duplicates were removed again with the same filtering procedure. We also remapped published ancient genomes following the same procedures for comparative analysis (Supplementary Table 3). Reads obtained from libraries were trimmed from both ends by 10 bps using *trimBAM* command of *bamUtil* software (*70*) to remove postmortem deamination artefacts.

#### Contamination estimates and genetic sex determination

Postmortem deamination patterns were estimated from pre-trimmed data using *PMDtools* (*71*) with the “*--deamination*” parameter. We conducted authenticity analysis using three additional approaches: *contamMix* (*72*) and *Schmutzi* (*73*), which make use of the rate of consensus mitochondrial sequence mismatches, and *ANGSD*, which estimates the excess of heterozygous positions for the haploid X-chromosome in male individuals (*74*). In order to detect non-endogenous reads, the *contamMix* library in R calculates a contamination probability using a reference panel of 311 diverse mitochondrial genomes. For this approach, consensus mitochondrial sequences were created using *ANGSD* (*74*) with parameters “*-doFasta 2 -doCounts 1 -minQ 30 -minMapQ 30 -setMinDepth 3 -r MT*”. *Schmutzi* also calculates probability of authenticity using deamination patterns on the consensus mitochondrial DNA fragments, but it additionally includes the information from read lengths and postmortem deamination since the longer and non-deaminated fragments are potential contamination sources (*73*). Finally, for the male individuals belonging to our sample set, contamination based on X-chromosome was estimated by running the *ANGSD* (*74*) algorithm with the command *‘‘angsd -i BAMFILE -r X:5000000 -154900000 -doCounts 1 -iCounts 1 -minMapQ 30 -minQ 30’’* for X-chromosome mapped reads. The probability of heterozygosity on the X-chromosome was then calculated using the Rscript *contamination.R* (*72*) and the reference files provided in the *ANGSD* package (*74*).

In order to determine the genetic sex of ancient individuals, we used the script *ry_compute.py* (*75*), which computes *R_Y_*, the ratio of the number of reads mapped to the Y-chromosome to the number of reads mapped to both the X- and Y-chromosomes per *BAM* file.

#### Uniparental markers

Y-chromosome haplogroups of male individuals were assigned using *PhyloTreeY* (*76*) (http://phylotree.org/Y/tree/). SNPs in the analysis were retrieved from the International Society of Genetic Genealogy (ISOGG) 2016 database (https://isogg.org) (v.11.04). The variants of the samples were called using *SAMtools* (v.1.9) *mpileup* with the “*–B*” parameter (*68*) and positions with base quality and mapping quality of less than 30 were filtered out.

Consensus mitochondrial sequences were produced from the sequence alignment files using *ANGSD* (*74*) with parameters “*-doFasta 2 -doCounts 1 -minQ 30 -minMapQ 30 -setMinDepth 3*”. Breadth of coverage was calculated for the positions which have depth ≥ 3 (Supplementary Table 2). Then, we identified haplogroups for individuals in which we recovered more than 15% of mitogenome by using *HaploGrep* (*77*).

#### SNP dataset preparation

After remapping previously published ancient genomes with the above-described procedure (see the section “Sequencing Data Preprocessing” above), SNP calling were done in two steps: i) *pileup* files were generated with the *SAMtools* (v.1.9) *mpileup* software (*78*), and ii) pseudo-haploid genotypes were generated by randomly choosing one allele per SNP locus from all *BAM* files using the *PileupCaller* v1.4.0.5 tool (https://github.com/stschiff/sequenceTools) and the option *“-randomHaploid”*. In this study, we used three different SNP panels.

1. <u>*Human Origins Dataset*</u>: Present-day Western Eurasian samples were extracted from the Human Origins dataset (*10, 25*) and merged with published Southwest Asian Early Holocene genomes (Supplementary Table 3) and the newly generated Çayönü samples. A total of 605,775 SNPs were included. This dataset was used to perform the principal components analysis.
2. <u>*1240K Dataset*</u>: We downloaded Allen Ancient DNA Resource (AADR) (https://reich.hms.harvard.edu/allen-ancient-dna-resource-aadr-downloadable-genotypes-present-day-and-ancient-dna-data) (v50.0) 1240K dataset and extracted Human Genome Diversity Project high-coverage genomes. After calling these positions in Southwest Asian Early Holocene genomes and Çayönü samples, we merged the datasets. The dataset consisted of 1,151,145 autosomal positions to compute runs of homozygosity.
3. <u>*Yoruba Dataset*</u>: We generated another SNP panel choosing bi-allelic sites with minor allele frequency (MAF) greater than 10% in the African Yoruba genome sample (56 female and 52 male, total 108 individuals) from the 1KG phase 3 dataset (1000 Genomes Project Consortium, 2015). We merged this with genotype files generated from high-coverage genomes (n=279) from the Simon Genome Diversity Project (SGDP) representing present-day global diversity (*80*). Additionally, we included published ancient genomes from Western Eurasia (Supplementary Table 3) and the Çayönü population to conduct population genetic analysis. After merging the datasets, remaining 5,991,735 autosomal SNPs in total were included. We also prepared another dataset including 220,384 X-chromosomal SNPs with the same procedure by removing the pseudoautosomal regions to conduct kinship analysis based on X-chromosome.

#### Genetic kinship analyses

In order to determine close kin up to third-degree relatedness, we estimated genetic kinship coefficients (θ) between each pair of individuals. To achieve this, we ran two alternative software, *READ* (*42*) and *NGSRelate* v2 (*41*), and jointly analysed their results. *READ* calculates pairwise mismatch rates (*P*_*0*_) of pseudo-haploid genome pairs in 1 Mbp windows and normalises these values with the median *P*_*0*_ of the sample, assuming the median represents a non-related pair (*42*). We used *1 - Normalized P*_0_ values as an estimate of the kinship coefficient (θ). *NGSRelate* v2 (*41*) calculates 9 Jaccard coefficients from genotype likelihoods, using background allele frequencies, and then computes θ = *J*_1_ + 0.5 × (*J*_3_ + *J*_5_ + *J*_7_) + 0.25 × *J*_8_. Since our Çayönü genome sample was composed of low coverage genomes, to increase the resolution of *NGSRelate*, we provided background allele frequencies calculated from a total of 211 Southwest Asia Holocene individuals including Çayönü, and published Anatolia, Levant, and Zagros genomes.

Of the 14 genomes produced, one pair (cay018 and cay020) were coherently identified as “Identical/Twin” by both methods. These two libraries were obtained from human remains from the same building and one library was constructed from a left-side and the other one a right-side petrous bone. The reconstructed skull showed that both petrouses could belong to the same infant. In addition, both libraries showed similar DNA preservation patterns (see Supplementary Table 2 for human proportions and damage patterns), which would be consistent with the possibility that the bones derived from a single individual. We thus merged these two libraries, leaving us with a total of 13 individuals.

After merging cay018 and cay020, we repeated the genetic kinship analysis with the 13 individuals but using autosomal and X chromosomal data separately. To increase confidence in the estimates, we stipulated a minimum number of overlapping SNP counts between pairs of individuals, >2000 SNPs when analysing autosomal data and >200 SNPs when analysing X-chromosomal data. We further required a significance of |Z| > 2 for the normalised *P*_*0*_ (estimated using variance among the genome-wide 1 Mbps windows by the *READ* software). Both *READ* and *NGSRelate* yielded well-correlated kinship coefficients (Spearman’s rho for autosomal θ = 0.65, for X-chromosomal θ = 0.49). One pair (cay004-cay033) was estimated as first-degree, two pairs (cay004-cay015 and cay011-cay027) as second-degree, and one pair (cay008-cay013) as third-degree kin according to autosomal kinship analysis. All these related pairs were co-buried in the same buildings. We excluded one of each pair of close kin (the lower coverage individual of each pair) from downstream population genetic analyses to ensure sample independence. Strikingly, cay008 and cay013 individuals were closer kin than third-degree according to X-chromosomal kinship analysis. Both individuals were females; while cay013 was an adult, cay008 individual was a 1.5-2 year-old child (Table S1).

To resolve the pedigree of the relationship between cay008 and cay013, we constructed and analysed possible maternal and paternal pedigrees of cay008 using the *pedsuite* package in R (*81*). Since she died before fertile age, cay008 cannot be the ancestor but could only be related to cay013 through her parents. Given this, we generated all possible third-degree relatives (great-grandparents, great-aunts, half-aunts and first cousins). Then, we calculated theoretical values of autosomal and X-chromosomal θ values for all pairs in the pedigrees (Fig. S5 and S6).

#### *f-* and *D-*statistics

We computed outgroup *f_3_*-statistics and *D*-statistics using the *qp3pop* and *qpDstat* programs, respectively, from the *ADMIXTOOLS* package (v7.0.2) (*25*) with default parameters. We used the present-day West African Yoruba sample (n=3) from the SGDP dataset (*80*) as an outgroup in both analyses. We ran pairwise outgroup *f*_*3*_-statistics for each individual from early Holocene Anatolia, Levant, Iran, and Upper Mesopotamia. We corrected *p*-values for multiple testing using the R *“p.adjust”* function’s false discovery rate (i.e. Benjamini-Hochberg) correction (*82*). In *D*-statistics involving Çayönü as a population, we performed analyses separately using the 9 genomes excluding one individual from close relatives and excluding cay008.

#### Isolation-by-distance analysis and within-population diversity

We computed geodesic geographic distance utilising the *geodist* package in R (*83*). To test isolation-by-distance, we computed the model fit with the *“lm”* function in R and compared geographic distance and genetic distance. We used 1-*f_3_* scores of each individual pair as a proxy for genetic distance. We filtered out pairs from the same site, pairs with >1000 years time difference between individuals, and pairs with <2000 overlapping SNPs.

To compare within-population genetic diversity of specific Neolithic settlements from Southwest Asia, we again utilised 1-*f_3_* scores between pairs of individuals as measure of genetic difference. We calculated significance using permutation tests through an in-house R script, where we randomised population identity with the R “sample” function.

#### Runs of Homozygosity Analysis

We used the Python package *hapROH* v.0.3a4 (https://pypi.org/project/hapROH/0.3a4/) to detect runs of homozygosity (ROH), which are long, homozygous stretches of the genome that result from the common ancestry of the maternal and paternal chromosomes (*34*). We used the default parameters of *hapROH* with pseudo-haploid genotypes, which contain more than 300,000 SNPs of the 1240K SNP Panel, the default genetic map of hapROH, and 5,008 global haplotypes from the 1000 Genomes Project (*33, 84*). We detected ROH for ancient genomes from Early Holocene Southwest Asia (Supplementary Table 3) and West and Central Eurasian present-day genomes (*85*). We performed linear regression using short ROH (4-8 cM) in present-day genomes to create a baseline that represents solely drift with no recent inbreeding. Right-shift from the baseline indicates that parents of this individual could be close-kin, whereas individuals that are around the baseline, and also have a relatively high number of ROH come from a population with potentially low *N_e_* (effective population size) (*34*). We filtered out ROHs <4 cM following the original *hapROH* publication which suggested that the method can detect ROH >4 cM (*33*).

### Dimensionality reduction analyses

#### Multidimensional scaling

We summarised the outgroup *f*_*3*_-statistics calculated across all pairs of individuals using multidimensional scaling and visualised the first two dimensions. First, we created a dissimilarity matrix of pairwise genetic difference (1-*f*_*3*_) values. From this we filtered out the pairs that had <2000 overlapping SNPs. We then applied the *“cmdscale”* function in R.

#### Principal component analysis

To perform principal component analysis (PCA), we utilised the *“smartpca”* (version 16000) software of *EIGENSOFT* (v7.2.1) (*86*) with the *“lsqproject: YES”* option to project ancient individuals onto principal components calculated on genome-wide polymorphism data of 55 Western Eurasian present-day populations (760 individuals) from the HumanOrigins SNP Panel (*10, 25*).

#### Admixture modelling

We modelled admixture proportions using the *“qpAdm”* software from the ADMIXTOOLS (v.7.0.2) package. We selected a reference differentially related to left populations covering modern and ancient diversity (*87*). We found that the following base reference set was able to distinguish our relevant populations: *Mbuti, Ust_Ishim, Kostenki14.SG, MA1, Han, Papuan, Dai, Chukchi, Mixe, CHG, Natufian, WHG, AfontovaGora3, Iberomaurusian*.

We then performed all possible two- and three-way models adding published genomes representing late Pleistocene and early Holocene populations of Central Anatolia, Zagros, and Levant as surrogates (“left populations”), and Çayönü genomes as targets. We ran all *“qpAdm”* analyses with *“allsnps: YES”* option, which is robust to low-coverage data (*87*). Any model without a Central Anatolia-related source did not work, yielding p-values < 0.05.

#### Visualisation

All plots were generated in R (*88*) using *ggplot* (*89*) and *ggpubr* (*90*) packages. Other packages used to analyse, clean and visualise the data are the following: *tidyverse* (*91*), *patchwork* (*92*), *reshape2* (*93*), *ggplotify* (*94*), *ggrepel* (*95*), *emojifont* (*96*), *ggforce* (*97*), *rgdal* (*98*), *raster* (*99*), *plyr* (*100*), *MetBrewer* (*101*), *pedsuite* (*102*).

## Acknowledgments

We thank all colleagues at the METU CompEvo and Hacettepe Human_G groups, the Diyarbakır Museum and the Ministry of Culture of Turkey for permissions to work on the material. The authors acknowledge support from the National Genomics Infrastructure in Stockholm funded by Science for Life Laboratory, the Knut and Alice Wallenberg Foundation and the Swedish Research Council, and SNIC/Uppsala Multidisciplinary Center for Advanced Computational Science for assistance with massively parallel sequencing and access to the UPPMAX computational infrastructure.

## Funding

European Research Council Consolidator Grant 772390 (MS)

## Author contributions

(a) N.E.A., A.Ö., Y.S.E., Ö.D.E., A.G., F.Ö., Ç.A., M.S. conceived and designed the study and experiments; (b) Ö.D.E., Y.S.E., A.Ö., S.S., M.M.K., G.Ç., provided and prepared the osteoarchaeological material; (c) A.Ö., S.S., H.C.G., C.K., Ç.A., compiled and analyzed archaeological data and building plans; (d) D.D.K. performed molecular biology laboratory experiments with support from V.K.L.; (f) N.E.A., A.A., D.K., K.B.V., E.Sa., K.G., M.Ö., analysed genetic data, supervised by F.Ö., A.Ö., M.S.; (h) N.E.A., A.Ö., Y.S.E., A.G., F.Ö., Ç.A., A.A., K.B.V., D.D.K., H.C.G., C.K., G.M.K., M.S., A.S., E.Sü., wrote the manuscript with contributions from all authors.

## Competing interests

Authors declare that they have no competing interests.

## Data and materials availability

The genomic alignment data (BAM format) is available through the European Nucleotide Archive (ENA) under the accession number XXX. All other data needed to evaluate the conclusions in the paper are present in the paper and the Supplementary Materials.

## Notes

### Competing Interest Statement

The authors have declared no competing interest.

