## Supplementary Figures and Table S1 for "A genomic snapshot of demographic and cultural dynamism in Upper Mesopotamia during the Neolithic Transition"

**This PDF file includes:**

Figs. S1 to S6
Table S1

**Fig. S1.**
Human proportion and coverage of the deep-sequenced specimen from the Çayönü population.

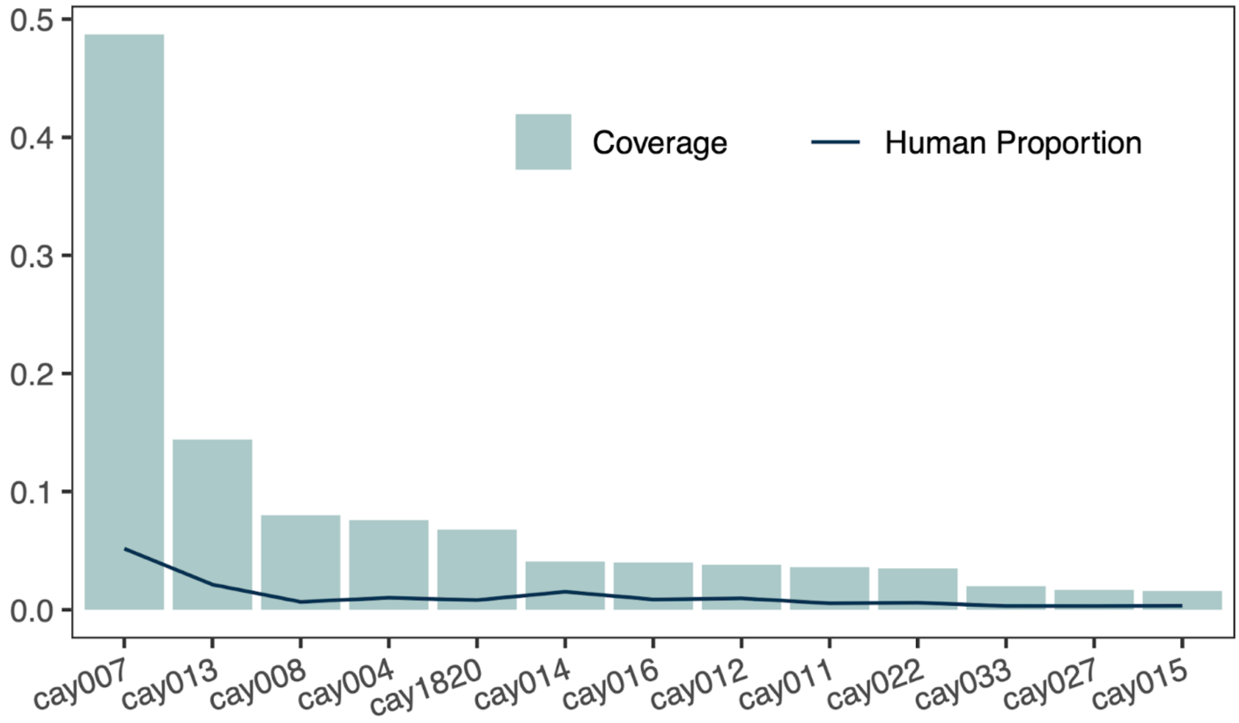

**Fig. S2.**
Comparison of the DNA preservation in Çayönü and contemporary Central Anatolia sites, Aşıklı and Boncuklu.

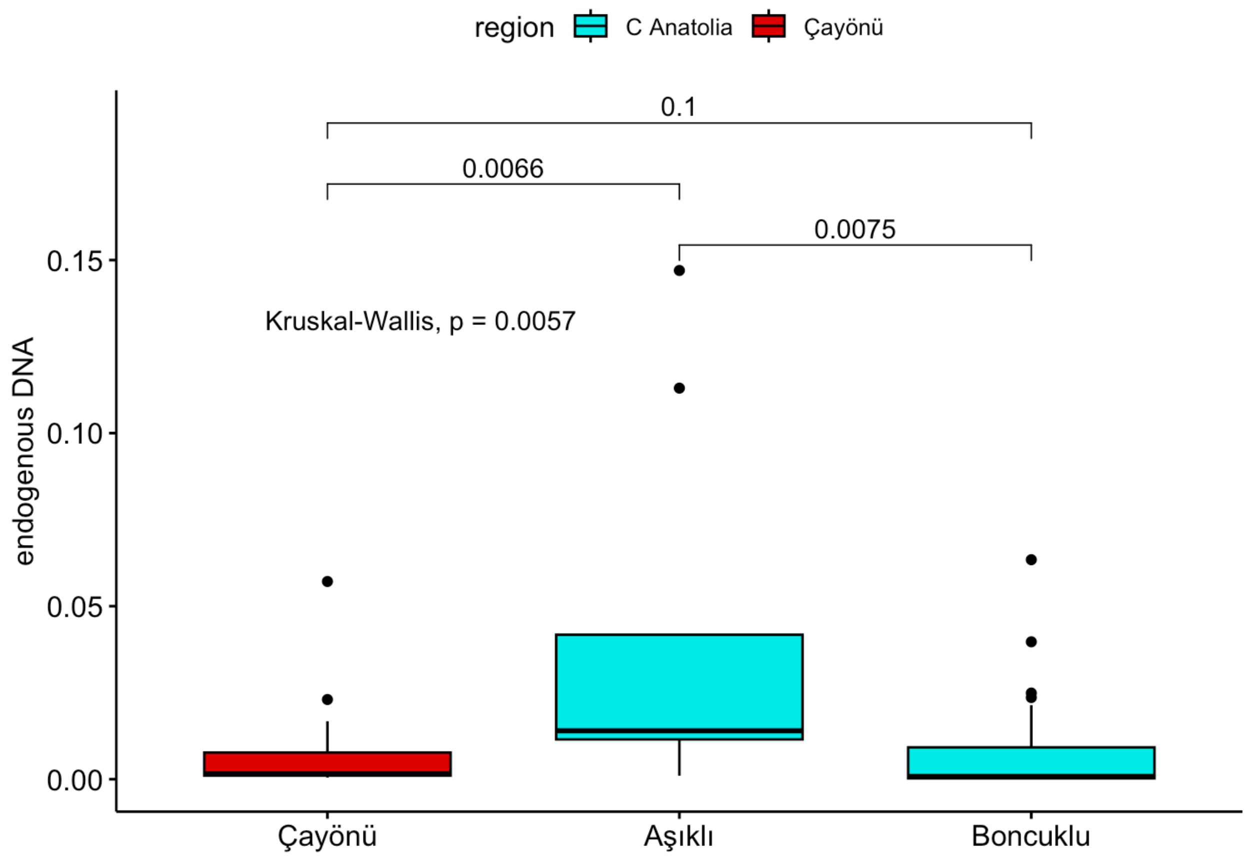

**Fig. S3.**

Principal Component Analysis (PCA) of Southwest Asian early Holocene populations which were projected onto 55 present-day West Eurasian populations. While a Southern Levant Neolithic individual (KFH2_KFH002) falls into the Anatolian cluster which is consistent with *(6)*, a Çayönü individual (cay015) with 6,986 SNPs appears in the Southern Levant cluster. This individual has the lowest coverage in our sample. The outlier individual (cay008) has been labelled in the figure.

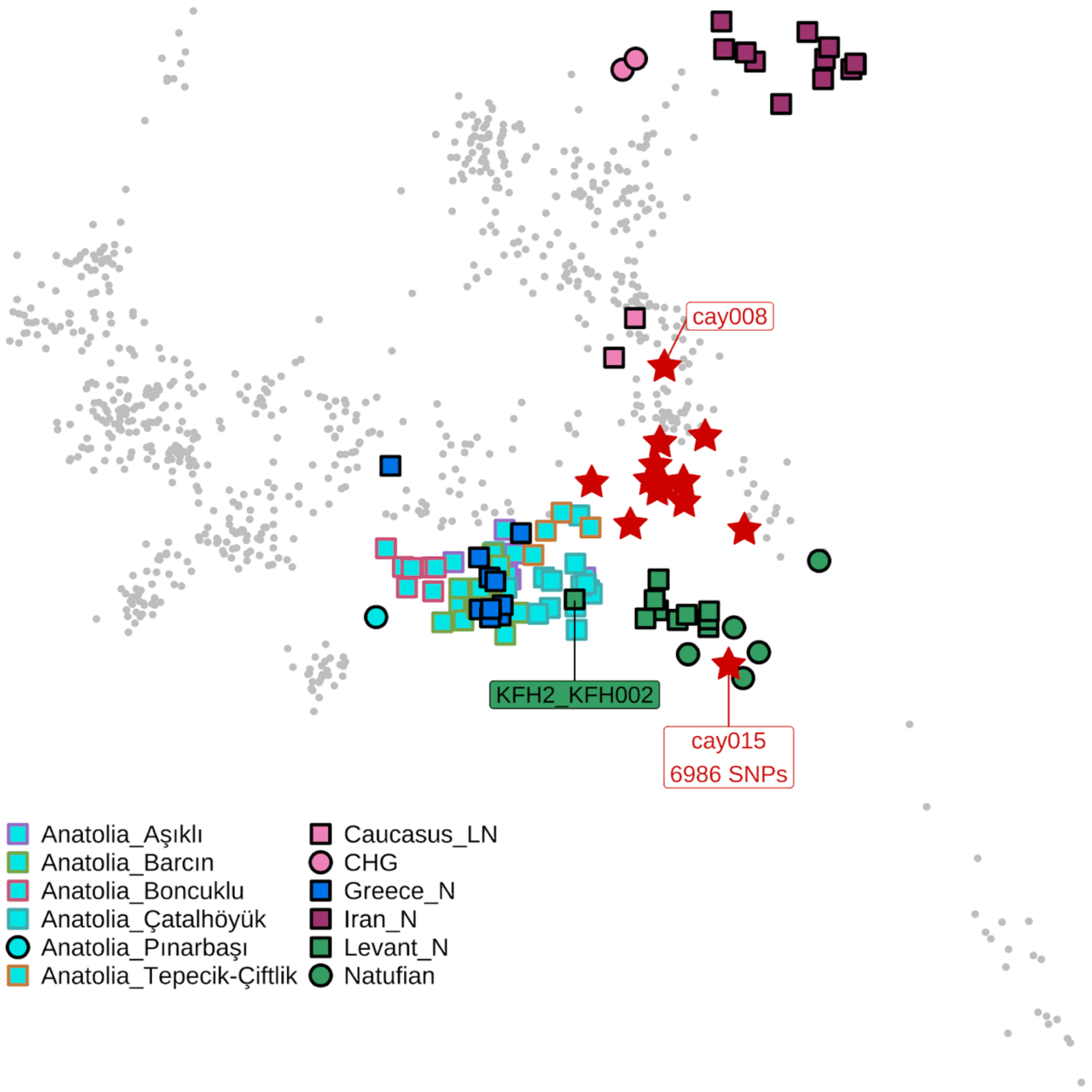

**Fig. S4.**

Comparison of genetic diversity (1 - $f_3$) of co-buried individuals in Central Anatolia and Çayönü.

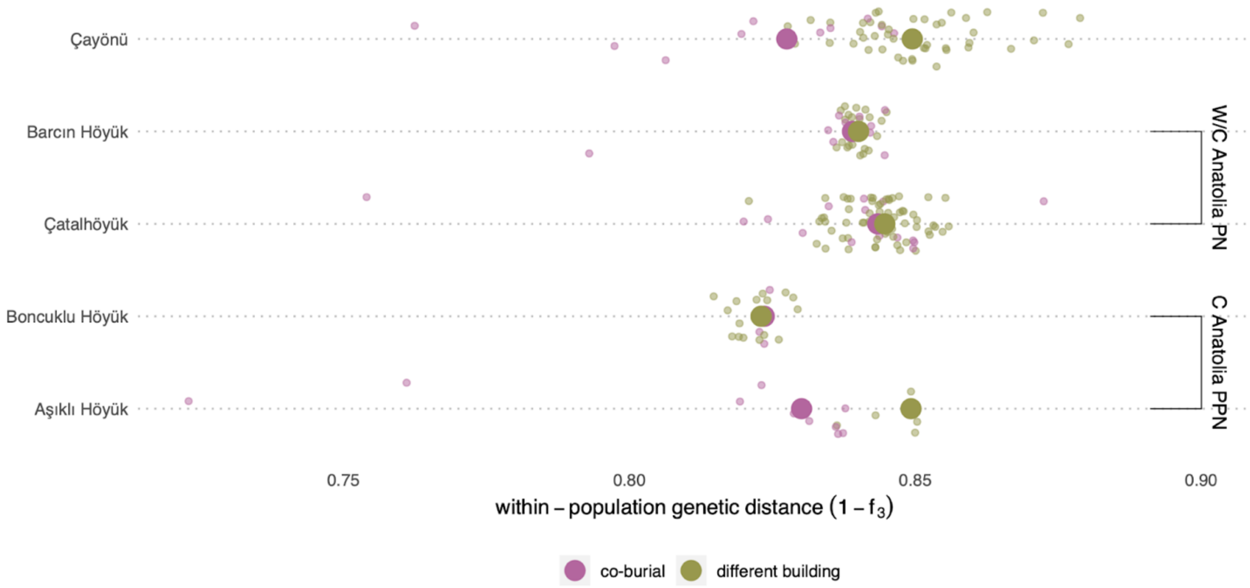

**Fig. S5.**

Simulated pedigrees for **(A)** paternal and **(B)** maternal relatives of cay008 to resolve her relationship with cay013 who is an adult female. Blue circles with dots correspond to the possible female third-degree relatives of cay008. Given that cay008 is an infant all relatives were simulated as related through the mother or father but not as descendants.

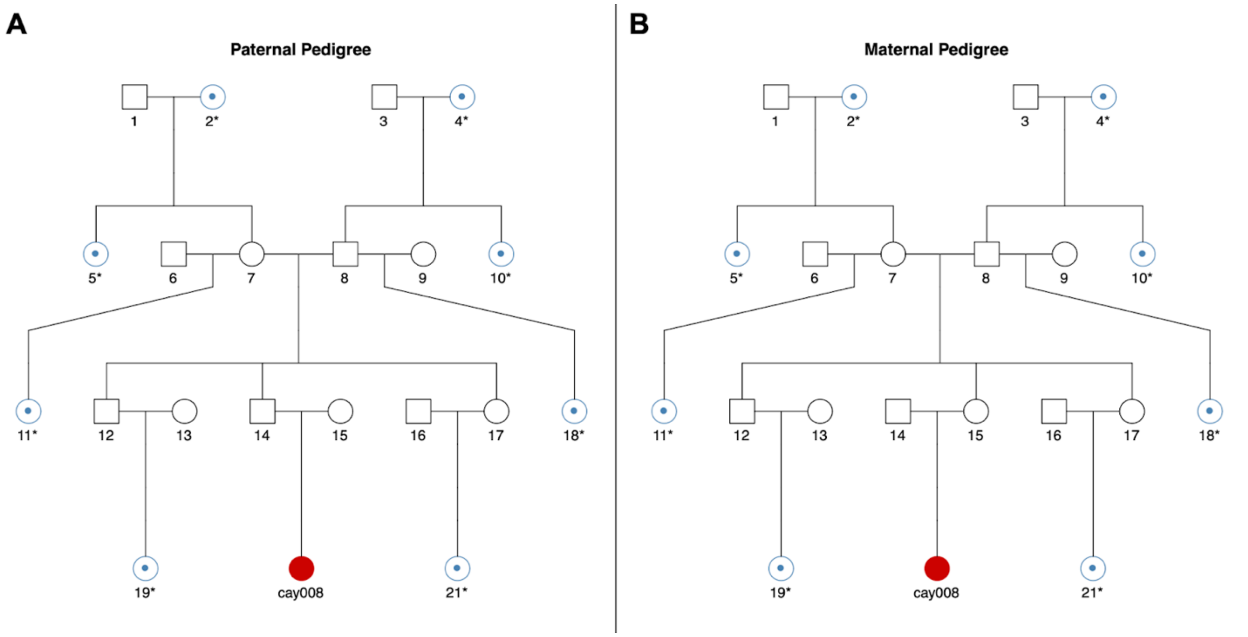

**Fig. S6.**

Theoretical kinship coefficient (θ) values between cay008 and her **(A)** paternal and **(B)** maternal relatives. Autosomal θ values are on the x-axes while X-chromosomal θ values are on the y-axes. The orange-shaded area shows potential values similar to real data where X-chromosomal θ is higher than autosomal θ. Red dots represent θ values *1 - Normalised* $P_0$ between cay008 and cay013 individuals, whereas horizontal and vertical bars infer 95% confidence intervals. Triangles show the potential female relatives falling in this range. Annotated numbers correspond to relatives in the two pedigrees shown in Supplementary Figure 5. Jitter was added to the points to visualize all the data.

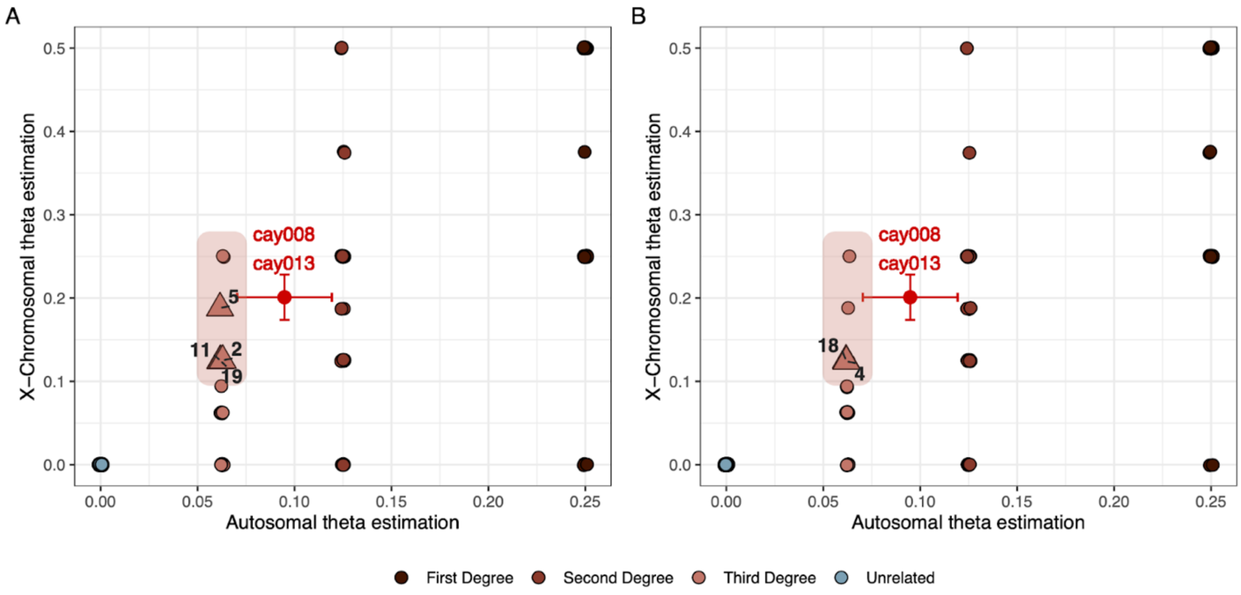

**Table S1.**

Archaeological contexts and anthropological characteristics of all screened Çayönü individuals.

| Building | Sub-phase | DNA lab No | DNA Preservation | Skeleton Number | Genetic Sex | Age at Death | Burial Type | Basic Pathologies |
|---|---|---|---|---|---|---|---|---|
| CA | c1 (PPNB) | cay001 | - | ÇT'86 S.1 | Male | 40-45 years old (Middle Adult) | Tightly flexed, NNE-SSW oriented, lying R, face down | Slightly developed osteoarthritis (OA) on the hip and shoulder joints; moderate arthritic changes in thoracic and lumbar vertebrae and knee joint. New bone formation on the internal table of the cranium and pelvic bones. |
| CA | c1 (PPNB) | cay002 | - | ÇT'78 S.12 | Unknown | 11-12 years old (Child) | Tightly flexed, primary, NE-SW oriented, lying R, face down | A healed depressed trauma on the right parietal bone; moderate cribra orbitalia on the right orbital roof. |
| CXa | c3 or c1 (PPNB) | cay003 | - | ÇT'72 S.10a | Female | 35-45 years old (Middle Adult) | Flexed, primary, W-E oriented, lying L, face S | A healed fracture on left big toe. |
| CL | c1(PPNB) | cay004 | + | ÇT'78 S.16 | Female | 33-46 years old (Middle adult) | Partially flexed W-E oriented, lying L, face SW | Healed fracture on the distal end of left ulnae; Monteggia fracture on right ulnae; periostitis on fibulae, slightly porotic hyperostosis on the cranium; slightly developed osteoarthritis on proximal end ulnae; Schmorl's nodule on 5th lumbar vertebrae. |
| CR | c3 (PPNB) | cay005 | - | ÇT'78 S.9 | Male | Adult (>15) | It was excavated between walls | Three healed depressed fractures (one of them is on the midline of the frontal bone and two of them are left parietal bone; slightly developed PH; severe osteoarthritic changes on the cervical vertebrae. |

| | | | | | | | | |
|---|---|---|---|---|---|---|---|---|
| CXa | c1(PPNB) | cay006 | - | ÇT'81 S.6 | Unknown | 4-5 years old (Child) | - | - |
| GE | Mid g (PPNA-PPNB transition) | cay007 | + | ÇT'81 S.2a | Male | 20-23 years old (Young adult) | Flexed, NW-SE oriented, lying R | A healed depressed trauma on right parietal bone and a healed fracture on felt 5th phalanges of the foot; slightly developed osteoarthritis on the right foot bones. |
| CA | c1 (PPNB) | cay008 | + | ÇT'86 S.2 | Female | 1-2 years old (Infant) | Flexed?, skull in W | Cauterization; circular type intentional head-shaping with post-coronal depression; woven bone formation on inner surface of the occipital bone. |
| CXa | c3 or c1 (PPNB) | cay009 | - | ÇT'72 S.8b | Female | 2-4 years old (Child) | Unknown, represented by teeth (she was found with ÇT'72 S.8a) | - |
| CN | c1 (PPNB) | cay010 | - | ÇT'78 S.2 | Unknown | 2-2,5 years old (Infant) | Flexed | Severe hypoplasia; Possible head-shaping with the sign of post-coronal depression. |
| CN | c1 (PPNB) | cay011 | + | ÇT'78 S.7 | Male | 33-42 years old (Middle adult) | Flexed, primary, SSE-NNW oriented, lying L, face down | Two healed depressed trauma (one of them is on left parietal and the other is on the frontal); healed periostitis on tibiae; Healed fracture on the left big toe, slightly osteoarthritis on the right ulnae and severe on cervical vertebrae. |
| CN | c1 (PPNB) | cay012 | + | ÇT'78 S.6 | Male | Nearly 12 years old (Child) | Tightly flexed, primary, SE-NW oriented, lying L, face down | Possible Scurvy? New bone formation is observed around the muscular attachment of all long bones and porosity on the ramus of the mandible. |

| | | | | | | | | |
|---|---|---|---|---|---|---|---|---|
| CA | c1 (PPNB) | cay013 | + | ÇT'78 S.21 | Female | 43-58 years old (Old adult) | Tightly flexed, NNE-SSW oriented, lying R, face down | Depressed trauma on the occipital; A fracture left costa, and periostitis on this fragment; slightly developed porotic hyperostosis and cribra orbitalia; slight osteoarthritis on the hand and foot phalanges; A possible head-shaping with a plano-occipital flattening. |
| CL | c1(PPNB) | cay014 | + | | Female | Adult | - | - |
| CL | c1(PPNB) | cay015 | + | ÇT'81 S.15 | Female | 40-45 years old (Middle adult) | Flexed, primary, SSW-N oriented, lying R, face W | A healed Colles' fracture on the left radius; a healed fracture on medial foot phalange; periostitis on sacrum, slight porotic hyperostosis and cribra orbitalia; slight osteoarthritis on the distal radius, thoracic and lumbar vertebrae. |
| CL | c1(PPNB) | cay016 | + | ÇT'81 S.8 | Female | 40-55 years old (Old adult) | Tightly flexed, S-N oriented, lying on the R | A healed Colles' fracture on the left radius; moderate porotic hyperostosis and cribra orbitalia; severe osteoarthritis on the left wrist bones, slightly on the right shoulder, moderate on the thoracic and severe on the lumbar vertebrae. |
| CV (CXa) | c3 or c1 (PPNB) | cay017 | - | ÇT'70 S.3 | Female | 28-37 years (Middle Adult) | Tightly flexed, S-N oriented, lying R, face SW | Cribra orbitalia on the left orbital roof. |
| GB(b) | g (upper) (early PPNB) | cay018 | + | ÇT'70 S.13 | Female | 3-9 months (Infant) | Flexed, partly twisted, primary, SW-ENE oriented, lying R | No pathology observed. |
| GB(b) | g (upper) (early PPNB) | cay019 | - | ÇT'70 S.10a | Female | Adult (>15) | Flexed, primary, ENE-WSW oriented, lying R | Healed depressed trauma on the R parietal bone; Hyperostosis frontalis interna; Disease idiopathic skeletal |

| | | | | | | | | |
|---|---|---|---|---|---|---|---|---|
| | | | | | | | | hyperostosis and intentional head-shaping with a post-coronal depression. |
| GB(b) | g (upper) (early PPNB) | cay020 | + | ÇT’70 S.11b | Female | 3-9 months (Infant) | Flexed, primary, lying R | Severe PH on parietal; infection on internal surface of occipital. |
| CXa | c1(PPNB) | cay021 | - | ÇT’72 S.9a | Female | Adult (>15) | Flexed, W-E oriented, lying R | Healed depressed trauma on the bregma region. |
| CR | c3 or cp3 (PPNB) | cay022 | + | ÇT’78 S.25 | Female | 9-12 months(Infant) | - | New bone formation on the internal table of the cranium; moderate porotic hyperostosis. |
| CA | c1 (PPNB) | cay023 | - | ÇT’72 S.5 | Female | 8-8,5 years (Child) | Flexed, secondary, NE-SW oriented, lying R, face E | Slight cribra orbitalia in both orbital roof. |
| CA | c1 (PPNB) | cay024 | - | ÇT’78 S.20 | Female | Middle adult | Tightly flexed, SE-NW orientation, lying L ?, face down | Slight porotic hyperostosis and cribra orbitalia. Osteoarthritic changes on cervical vertebrae. |
| CA | c1 (PPNB) | cay025 | - | ÇT’78 S.13 | Male | Adult (>15) | Tightly flexed, NNW-SSE oriented, lying R, face down | Healed fracture on the right 2nd and 3th metacarpal and distal phalanges, slight osteoarthritis on the right distal radius. |
| 21N | c2 (PPNB) | cay026 | - | ÇT’78 S.28b | Female | Old adult | - | Two healed depression trauma on the cranium (one is on the left frontal tuber and other is on the occipital; hyperostosis frontalis interna; slight osteoarthritis on right shoulder; moderate on the cervical vertebrae. |
| CN | c1(PPNB) | cay027 | + | ÇT’78 S.1 | Female | Adult (>15) | Flexed? secondary? SW-NE oriented, face? | Infection on the right tibia. |

| | | | | | | | | |
|---|---|---|---|---|---|---|---|---|
| CL | c1 (PPNB) | cay028 | - | ÇT'78 S.17 | Female | 22-24 years (Young adult) | Semi flexed, N-S directed lying on right side and partially on stomach | Slight porotic hyperostosis. |
| CA | c1 (PPNB) | cay029 | - | ÇT'78 S.19 | Female | 40-45 years (Middle adult) | Tightly flexed, SSW-NNE oriented, lying on the left side | Healed depressed trauma on the skull, maxillary sinusitis; periostitis on the right tibia; slight osteoarthritis on cervical vertebrae. |
| CXa | c1 (PPNB) | cay030 | - | ÇT'81 S.7 | Female | Adult (>15) | Flexed | Healed depressed trauma on the right parietal. |
| CXa | c1 (PPNB) | cay031 | - | ÇT'81 S. 4 | Male | Adult (>15) | Flexed | Two healed depressed trauma on the right parietal bone and slight porotic hyperostosis. |
| GE | Mid g (PPNA-PPNB transition) | cay032 | - | ÇT'81 S.2b | Male | Young adult | Together with 81 S.2a | Two healed fractures on the right foot bones; periostitis on the right tibia. |
| CL | c1 (PPNB) | cay033 | + | ÇT'84 S.60 | Male | 8-8,5 years (Child) | Under 81 S.15 (cay016) | Hematoma on the left femoral diaphysis. |
